# Combined impacts of climate and land use change and the future restructuring of Neotropical bat biodiversity

**DOI:** 10.1101/2021.07.23.453383

**Authors:** Fernando Gonçalves, Lilian P. Sales, Mauro Galetti, Mathias M. Pires

## Abstract

Forecasting the effects of global change on biodiversity is necessary to anticipate the threats operating at different scales in space and time. Climate change may create unsuitable environmental conditions, forcing species to move to persist. However, land-use changes create barriers that limit the access of some species to future available habitats. Here, we project the impacts of climate and land-use change on 228 Neotropical bat species by forecasting changes in environmental suitability, while accounting for the effect of habitat type specialization and simulating dispersal across suitable patches. We also identify the most vulnerable ecoregions and those that may offer future stable refugia. We further investigate potential functional changes by analysing the response of different trophic guilds. We found that the range contraction of habitat specialists, especially frugivores, was more frequent and stronger under all simulated scenarios. Projected changes differ markedly across ecoregions. While the Amazon region is likely to undergo high turnover rates in bat composition, the Andean grassland, Cerrado and Chaco might experience the greatest losses. The expansion of habitat generalists, which forage in open areas and commonly establish large colonies in manmade structures, coupled with the range contraction of habitat specialists is projected to homogenize bat communities across the Neotropics. Overall, dispersal will likely be the key for the future of Neotropical bat diversity. Therefore, safeguarding the refugia highlighted here, by expanding and connecting the existing network of protected areas, for example, may allow species to move in response to global change.

## Introduction

Climate and land use change are among the main processes driving biodiversity loss in the 21^st^ century (IPCC, 2019). Vulnerability assessments suggest that the Neotropics are where the combined impact of climate change and habitat loss will be the greatest (Colwell et al., 2008). According to these projections, most of the Neotropical region will be subjected to novel climates, while increasing rates of land use change will reduce habitat availability even further (Staude et al., 2020). Species’ responses to global changes depend on extrinsic and intrinsic factors related to the rate and magnitude of change in environments, as well as the species’ autecology and climatic tolerances. Exposure to non-analog climates may lead to physiological stress with likely reductions in individual fitness and thus population persistence in the long term (Ribeiro et al., 2016). Furthermore, if species are not able to adapt *in situ* or to track suitable climates, population declines and local extinctions may result in range contractions (Zamora-Gutierrez et al., 2018).

Land-use change represents an additional challenge for species dealing with climate change, as human-modified landscapes obstruct species movements by converting habitats into dispersal barriers (Sales et al., 2019). In the Neotropics, where deforestation and climate change pose huge threats to biodiversity, the combination of extrinsic and intrinsic stressors is expected to drive the decline of habitat specialists, which have limited diets and specific environmental requirements to roost and forage (Devictor et al., 2008; Clavel et al., 2011). Conversely, habitat generalists that thrive in a wide variety of environmental conditions and use a variety of resources, may even increase in abundance and geographic distribution in response to changes in the environment (Gonçalves et al., 2017; Farneda et al., 2019). Thus, advancing our understanding of the effects of environmental change on biodiversity requires examining how threats may impact species differently according to their ecological characteristics as well as identifying vulnerable and important conservation areas (Sales et al., 2019).

The threats related to the changing climate and habitat loss extend to virtually all clades, and impacts can already be observed, resulting in reduced distribution in invertebrates (Larsen, 2012) and vertebrates (Tingley et al., 2009; Sinervo et al., 2010). Because of their prompt response in terms of richness, abundance and physiology to changes in land use, management intensities and extreme weather events, bats are regarded as bioindicators that help understanding the impacts of environmental change in a broader context, (Sherwin et al., 2013; Gonçalves et al., 2017). Bats can be sensitive to overheating (Crawford and O’keefe, 2021), and changes in distribution of multiple species seem to have happened as a consequence of recent climate change (Wu, 2016). Certain species may even benefit from climate changes, expanding their ranges to regions that are currently unfavourable. For example, in Costa Rica many bat species have recently been documented at higher-than-normal elevations, suggesting range expansions (LaVal, 2004; Arias-Aguilar et al., 2020). Local extinctions and colonization dynamics in response to climate forecasted to the 21^st^ century may redistribute bat species (Pearson and Dawson, 2003; Zu 2015). Yet, changes in land use could limit populations in tracking their bioclimatic niches (Mendes et al., 2021) Thus, forecasts of the effects of environmental change on Neotropical bats are important not only to understand how distribution patterns in this diverse group may be altered, but also for a broader assessment of how climate and land use will reorganize biodiversity.

Here, we infer the combined impacts of climate and land use change on Neotropical bats focusing on potential changes in spatial patterns of bat richness. Using species distribution models we generate projections of changes in environmental suitability for a representative sample of the Neotropical bat biota (ca. 80% of all species) under two (Mitigation A1B and Business-as-usual A2) climate and land use socio-economic development scenarios for the end of the 21^st^ century. We also investigate three different dispersal scenarios to test how important natural and anthropogenic barriers can be in determining future spatial distribution patterns. Based on the resulting projections we examine how differences in habitat use and diet may affect distribution patterns and, thus, reshape the configuration of bat diversity across Neotropical ecoregions.

## Methods

### Study system

To assess the effects of climate and land use change on Neotropical bat distributions we studied the distribution of 228 Neotropical bat species, which corresponds to ca. 80% of documented species in the region. The species not included in the study are species for which there is limited information on distribution or habitat use or those whose distribution is too restricted (see Appendix S1 for further details on the Methods and Appendix S2 for more information on the studied species). We focused our analyses on the Neotropics, ranging from northern Mexico to central Argentina and including the whole Caribbean (Morrone 2014). We also investigated differences across Neotropical ecoregions by adopting the Neotropical division into 10 ecoregions proposed by Antonelli et al. (2018): Amazonia, Atlantic Forest, Andean Grassland, Cerrado and Chaco, Caatinga, Dry Western, Dry Northern, West Indies and Mesoamerica (Fig. 1).

**FIG. 1:**
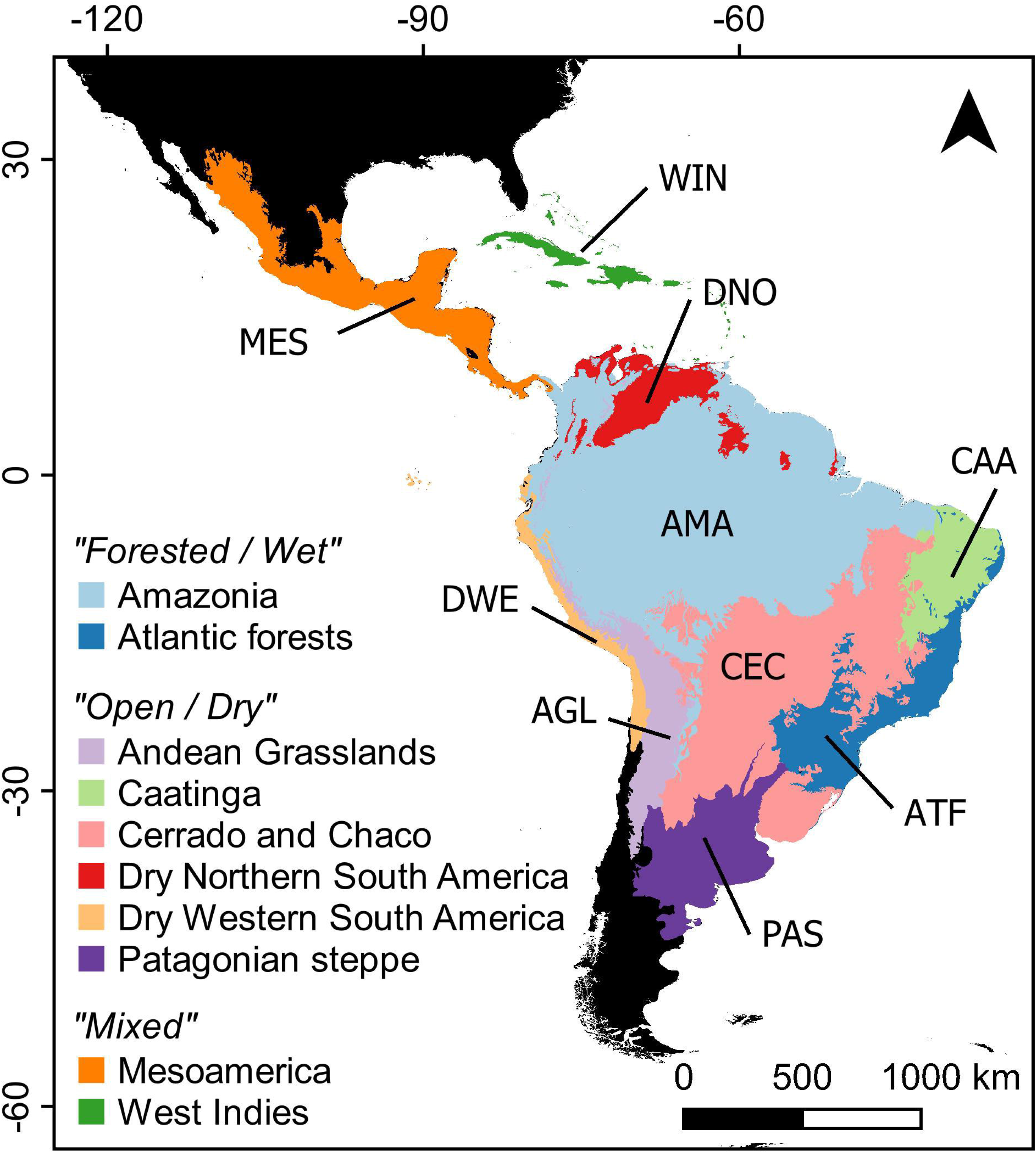
Neotropical region limits (sensu Olson et al., 2001) and ecoregion classification (sensu Antonelli et al., 2018) adopted in the analyses.

### Ecological niche models

To assess potential changes in species distribution patterns, we fitted ecological niche models calibrated with climate data from locations within which species are known to occur. Ecological niche models are correlational procedures aimed at assessing the environmental conditions associated to the occurrence of species in order to estimate their potential distribution, i.e. the locations where the environment is similar to that within the species’ known distribution. We used the species ranges available in the IUCN database (www.iucnredlist.org, date of access: August 08th, 2020) as the source for distribution information. For each species, we sampled a number of random points within the species inferred range that was proportional to its range size and used species distribution models to generate potential distribution maps. To do so, we first rasterized the IUCN range maps into a gridded file of 10’ resolution (approximately 0.17 degree of lat/long) and, to be conservative and avoid overestimation, we restricted our analysis to species whose range size was larger than 30 cells. For species whose range size was larger than 1000 cells, we sampled 12.5% of these cells, thus considered “presences” in the rasterized map. For species with range size varying from 501 to 1000 and 101 to 500 we sampled 25% and 50% of the cells within species ranges. Finally, for species with less than 100 cells within the total range we used all cells as occurrences to calibrate distribution models. Because we used IUCN range maps as a source of distribution information, the outcome of our ecological niche models represents the environmental conditions most frequently observed across species’ known distributional limits. Therefore, our results should not be interpreted in terms of probability of occurrence *per se*, neither can be directly translated in any abundance-related metric. Such broadly defined climate envelopes are meant to provide an initial assessment of species climatic suitability at the continental scale and are useful to investigate macroecological relationships between biotas and environments (Sales et al., 2020; Sales et al., 2020a, Sales et al., 2020b). Therefore, we stress that our results should not be considered at face value in conservation assessments at local spatial scale.

We acknowledge that using information from range maps to assess species climatic niches is not the “gold-standard” choice in ecological niche modelling, and that a comprehensive and non-autocorrelated dataset taken from confirmed on-ground presences and absences would be ideal (Araújo et al., 2019). However, this scenario is impractical for multispecies studies at continental scale, especially for large extensions of biodiversity-rich yet remote and under-sampled locations, such as most of the Neotropics (Etard et al., 2020; Zizka et al., 2021). Moreover, capture rates of bats are subjected to the complexity of the environment, being generally higher in open environment than in habitat with a more complex vegetation structure like forests. Although this source of bias may also affect range maps, the impact is much greater for the distribution of occurrence points. Data gaps (e.g. tropical rainforest and mountain top regions) in the distribution of Neotropical bats limit our ability to conduct large-scale analyses based on occurrence points, whereas biases can impact the validity of extrapolations (Aguiar et al., 2020; Delgado-Jaramillo et al., 2020). Furthermore, the sub-setting of realized niches due to defaunation and other non-climatic stressors creates truncated relationships between environmental suitability and environmental variables, underestimating biodiversity predictions (Faurby and Araújo, 2018) and overestimating the impact of future stressors (Lima-Ribeiro et al., 2017). Thus, by using species’ range maps the basis for distribution models instead of occurrence points with critical gaps and biases, we produce a conservative assessment and our analyses should be interpreted as a baseline of the potential effects of climate change on bat distribution.

To build distribution models we obtained climate information as 10’ resolution gridded raster files for land areas, derived from interpolation of worldwide ground weather stations, available in CliMond (https://www.climond.org/). Date referred to “present-day” conditions encompass averages of measurements between years 1961 – 1990. Although CliMond data includes 35 bioclimatic predictors, we only used a subset of predictors when calibrating ecological niche models. Before selecting among predictors, we built species-specific study areas, limiting the background extent of each species to regions likely accessible via migration (Barve et al., 2011). To do so, we defined a bounding box to extreme latitude/longitude coordinates, plus an additional 10 degrees to each bound and cropped climatic layers to adjust to the study extent. Then, we extracted the climatic information related to each species background extent. To select among predictors, we used a hybrid approach that combines pairwise correlations with variance inflation factor (VIF) (Marquardt, 1970), implemented in *usdm* R package (Naimi et al., 2014; R Core Team, 2021). After measuring the correlation between pairs of climatic variables, the pair with highest correlation value (threshold = 0.6) was identified and the variable with higher VIF was excluded from the pair. This procedure was repeated until all the strongly correlated variables were excluded.

After defining a proper set of uncorrelated predictors for each species models, we used an ensemble-based ecological niche modelling approach, using the *usdm* R package (Naimi et al., 2014). To accommodate the sensitivity of our predictions to method variation, we used two algorithms: boosted regression trees (BRT) and maximum likelihood (MaxLike). While boosted regression trees join recursing binary to adaptive boosting, combining several models in an ensembled prediction with higher performance (Elith and Leathwick, 2009), MaxLike is a method based on maximum likelihood, whose parameter output is formally related with the widely-used Maxent software (Merow and Silander, 2014). Both methods rely on machine-learning approaches, which are known to accurately predict simulated response curves (Elith and Leathwick, 2009), notably in cases of small sample size (Pearson et al., 2007).

The outcome of our ecological niche models was, then, evaluated for their accuracy using a repeated sub-sampling procedure where 25% of the records were regarded as the “test” sub-set, used to measure the performance of the models that were fitted using the remaining 75% of the records, the “training” sub-set. This procedure was repeated 100 times for each modelling method *per* species. We used the area under the receiver operating characteristic curve (AUC), a threshold-independent metric (Fielding and Bell, 1997), to evaluate the accuracy of our predictions (Appendix S2). AUC values of model performance vary from 0 to 1, where a value of 0.5 indicates accuracy similar to that of a random model a value of 1 indicates a perfect discrimination by both metrics. Because binary predictions were required as input in our simulation of dispersal via cellular automata (see below), we also calculated the true skill statistic (TSS) of the models to determine proper thresholds for each species and then created binary maps of suitable/unsuitable cells.

### Climate change forecasts

To generate projections of the potential effects of climate change on the potential distribution of Neotropical bats, we projected ecological niche models according to forecasts of climate dated to years 2030, 2050, 2070 and 2090. The fact that we project changes for every 20 years allowed us to track climatic niche bridges between successive timesteps (Littlefield et al., 2017). Accommodating the inherent uncertainty of climate change forecasts, we used two possible future scenarios, based on different expectations of human population growth, socioeconomic development, and associated emissions of greenhouse gases. The *Mitigation* scenario, A1B, estimates a temperature increase of 1.7 – 4.4°C by the end of the 21^st^ century, whereas the *Business-as-usual*, A2, forecasts a 2.0 – 5.5°C temperature increase. Those forecasts roughly correspond to the representative concentration pathways *RCP6.0* and *RCP8.5* from the IPCC-AR5 (IPCC, 2014) and project temperature increases above the 1.5°C threshold recommended to avoid the deleterious effects of climate change on ecosystems and human well-being (IPCC, 2019) but are congruent with observed changes from years 2000-2010 (IPCC, 2014) (see Appendix S1 climate data for details).

### Habitat filter masks

Ecological niche models calibrated with IUCN range maps are based on the assumption that any species can occupy all of the habitat within its extent of occurrence, which turns predictions overly “optimistic” and, thus, subjected to commission error (Lobo et al., 2010). To minimize these erroneous relationships in our maps of potential distribution, we created species-specific land cover masks, based on the IUCN habitat classification scheme, the most comprehensive effort available to characterize species habitat affiliations (https://www.iucnredlist.org/resources/habitat-classification-scheme). The major habitat types according to this scheme are forest, savanna, shrubland, grassland, wetlands (inland), rocky areas (e.g., inland cliffs, mountain peaks), caves and subterranean (non-aquatic), arid or semi-arid (desert), artificial-terrestrial (e.g. Pastureland, Plantations, Urban areas and heavily degraded forest). We considered habitat specialist species all those species whose occurrence was restricted to a single major land cover type (e.g. forest), regardless the specific category of land cover (88 species). Accordingly, we consider a habitat generalist species any species whose occurrence is not restricted to a single major land cover type (140 species). Species were, assumed to be unable to occupy major habitat types where they had not been recorded before. As these habitat types can be matched with terrestrial land cover, known as the area of habitat (Brooks et al., 2019), we grouped the habitat data into six major land cover classes (forest, savanna, grassland, farmland, urban, and barren). By doing so, we were able to reconcile the information on species-habitat associations to a global model that project changes on land-use and land-cover (LULC) (Li et al. 2017, described in Appendix S1 land-use and land-cover section).

### Dispersal assumptions

In heterogeneous landscapes, the quality and permeability of the matrix act as environmental filters that affect species’ persistence and ability to colonize newly suitable areas (Gonçalves et al., 2017; Farneda et al., 2019). Here, we explicitly incorporate dispersal into the future estimates of potential distribution assuming that expansion of a species distribution depends on the availability of habitat that allows dispersal across space. Specifically, we consider that dispersal is more likely across land cover types corresponding to habitat types where a species is known to occur, whereas land cover types that are not listed as major habitat types or habitats used for that species are considered barriers to movement across suitable climatic pockets. Therefore, we assume that dispersal limitation can prevent a species to access and occupy suitable environments in the future (see Appendix S3). To simulate dispersal we used the R package MigClim (Engler et al., 2012), a cellular automata model where dispersal depends on landscape barriers and species-specific dispersal limitations contingent on environmental suitability. We used the following parameterization of function “MigClim.migrate” for all dispersal simulations. The function argument hsMap included the habitat suitability models for intermediate timesteps (years 2030, 2050 and 2070); envChgSteps and dispSteps, i.e. the number of environmental change steps and the number of dispersal steps were set to three (the number of intermediate timesteps). The argument barrier included a raster file with the land-use and land-cover types listed as unsuitable for each species, and barrierType was defined as “weak” or “strong”, depending on habitat generalization or specialization, respectively. Finally, replicate NB was set to three full simulations of dispersal per species. For the remaining arguments we used the function default values (for further details on the dispersal simulation see the section modelling occupancy dynamics and dispersal in Appendix S1). To understand how dispersal constraints may affect future climate-driven range dynamics, we examined future patterns of potential distribution considering three different assumptions on dispersal ability. The 1) *Unlimited Dispersal* scenario, where bat species are assumed to have the ability to disperse across any type of land cover and fragmented landscape. The 2) *No Dispersal* scenario, where species are not allowed to move outside current distribution limits and estimates of change in climatically suitable areas are restricted to grids within present-day extent of occurrence. The (3) *Limited Dispersal* scenario, which is more realistic than the previous scenarios and assumes that bat species have the ability to disperse across areas where land cover is analogue to that where it occurs in the present, whereas no-analog environment is considered a barrier to dispersal. By performing simulations under these three different scenarios, we intended to infer the roles of environmental suitability and accessibility on the projected outcomes

For each combination of climate change, land use change and dispersal limitation scenarios we registered whether the potential range of each species would decrease or increase in the future. Based on the projected ranges we estimated the number of species that could potentially occupy each cell for the Neotropics as a whole and for each ecoregion (see Appendix S1 - modelling occupancy dynamics and dispersal). To understand how the reorganization of bat assemblages may translate into shifts in ecological functions, we also classify species according to trophic guilds and analyse the changes in spatial patterns for each guild. For the analysis of which trophic guild might be affected, we classified bats based on their main food items (feeding habit) according to Simmons (2005): Phytophagous bats: (1) frugivore and (2) nectarivore; Animalivorous bats: (3) carnivores, (4) sanguivores, (5) insectivores and (6) piscivores; and species that feeding on both animal and plant: (7) Omnivore (for further details about models and dispersal scenarios, see Appendix S1-modelling occupancy dynamics and dispersal).

## Results

### The magnitude of range shift

The magnitude of range shift in response to climate and land use change varied considerably according to different scenarios of climate change and dispersal limitation (Fig. 2). Simulating a realistic dispersal scenario, in which dispersal across space is limited by barriers of non-analog environment, most of the habitat-generalist species could expand their ranges in both (A1B) the Mitigation (A1B_LD_G*_median_* = 26 ± 1374%) and (A2) B.A.U. scenario (A2_LD_G*_median_* = 4 ± 1148%), but most habitat-specialist species are projected to experience major reductions in their ranges in the A2 scenario (A1B_LD_S*_median_* = 32 ± 347% and A2_LD_S*_median_* = −47 ± 217%) (Fig. 2 and Appendix S2).

**FIG. 2.**
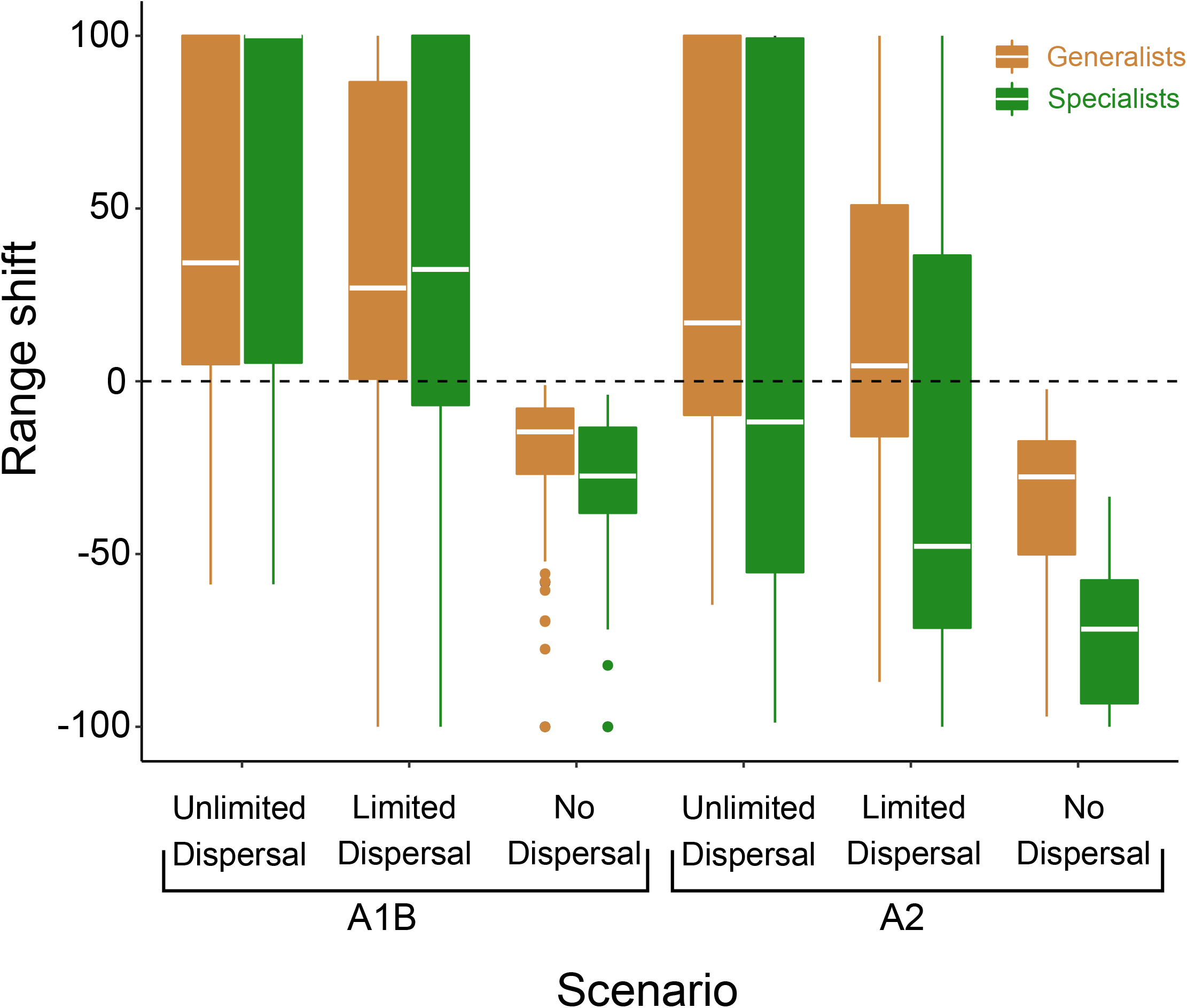
Proportional change in potential distribution for 140 habitat-generalist bat species and 88 habitat-specialist bat species in response to combined effects of climate and land use change. Projections correspond to two different scenarios for climate/land-use changes and different levels of dispersal limitation. A1B and A2 refer to Mitigation and B.A.U., i.e. more optimistic and less optimistic climate and land use change scenarios. Unlimited Dispersal assumes that bat species have ability to disperse across any type of environment and across fragmented landscape. Limited dispersal assumes that bat species have the ability to disperse only across analog environment, whereas non-analog environment is considered a dispersal barrier. In the No Dispersal scenario species are restricted to their current range.

Next, we simulated two dispersal scenarios representing opposite extremes regarding dispersal to test how accessibility changes the outcomes of environmental changes for Neotropical bats. Assuming that bats are confined to their current distribution limits, even the generalist species regarding land cover would be expected to lose a large proportion of their potential distribution (A1B_ND_G*_mean_* = −14 ± 18% and A2_ND_G*_mean_* = −27 ± 22%). Again, habitat-specialist species would be more impacted showing largest range reductions (A1B_ND_S*_mean_* = −27 ± 19% and A2_ND_S*_mean_* = −71 ± 19%; Fig. 2). This suggests that the area that is climatically suitable within the distributions of all species will reduce in the future.

Under the *Unlimited Dispersal* scenario, in which bat species have ability to disperse through any type of matrix, most of the habitat-generalist species could expand their ranges in both climate change scenarios (A1B_UD_G*_median_* = 34 ± 2437% and A2_UD_G*_median_* = 16% ± 2236%), but most habitat-specialist species are still projected to undergo range contraction in the A2 scenario (A1B_UD_S*_median_* = 122 ± 559% and A2_UD_S*_median_* = −11% ± 217%) (Fig. 2). This suggests that although within current ranges suitability will decrease, the total area of suitable climate will increase for most species.

### Neotropical ecoregions with higher projected impact

Despite projected losses in richness in many areas (Table 1), the overall outcome of projected range shifts was an increase in the mean richness at the local (cell) scale across most ecoregions (Fig. 3; Table 2), owing to the range expansion of habitat-generalist species (Fig. 4). However, at the regional scale, most ecoregions are projected to undergo a net loss in species richness (Table 1; Fig. 4). The Amazon was the region projected to experience the largest gains in richness at the local (cell) scale (Fig. 3; Table 2), especially nearby Andean mountains and Guiana Highlands (Fig. 4). Yet, these range shifts resulted in no net gains or losses in richness at the regional scale (Table 1). The Dry Western South America is the region projected to experience the greatest proportional gains, with richness increasing more than 2.5 times at the local scale (from 4 to 11 species; Fig. 3; Table 2) and 61 new species at the regional level (Table 1). Despite little change in local richness, Mesoamerica and the Patagonian steppe are also projected to gain species, with 28 and 10 novel species, respectively (Table 1). In contrast, the Andean grassland is projected to lose 22 species at the regional level in spite of a substantial increase in local richness (from 3 to 10 species) (Table 1). All remaining ecoregions are projected to suffer more loses than gains with the Caatinga, the Cerrado/Chaco, and the West Indies losing more than 15% of their current species (Table 1).

**Table 1:** Potential richness variation of Neotropical bat species at regional scale under the business-as-usual (B.A.U) scenario of climate and land-use changes and assuming limited dispersal (bat species have the ability to disperse only over analog environment).

| Ecoregions | Present | Future |  |  |  |
| --- | --- | --- | --- | --- | --- |
|  | Species richness | Species gains | Regional extinction | Global extinction | Species richness |
| **<u>Forested Wet</u>** |  |  |  |  |  |
| Amazonia | 212 | 0 | 0 | 0 | 212 |
| Atlantic forests | 98 | 3 | 13 | *D. capixaba/L. mordax* | 86 |
| **<u>Open Dry</u>** |  |  |  |  |  |
| Andean Grassland | 160 | 8 | 21 | 0 | 147 |
| Caatinga | 79 | 6 | 17 | 0 | 68 |
| Cerrado and Chaco | 140 | 14 | 21 | 0 | 133 |
| Dry Northern | 172 | 3 | 14 | *Phyllostomus latifolius* | 161 |
| Dry Western | 103 | 61 | 7 | *Eptesicus innoxius* | 157 |
| Patagonian steppe | 30 | 10 | 7 | 0 | 33 |
| **<u>Mixed</u>** |  |  |  |  |  |
| Mesoamerica | 105 | 28 | 0 | 0 | 133 |
| West indies | 27 | 0 | 6 | *Chilonatalus micropus* | 21 |

**FIG. 3.**
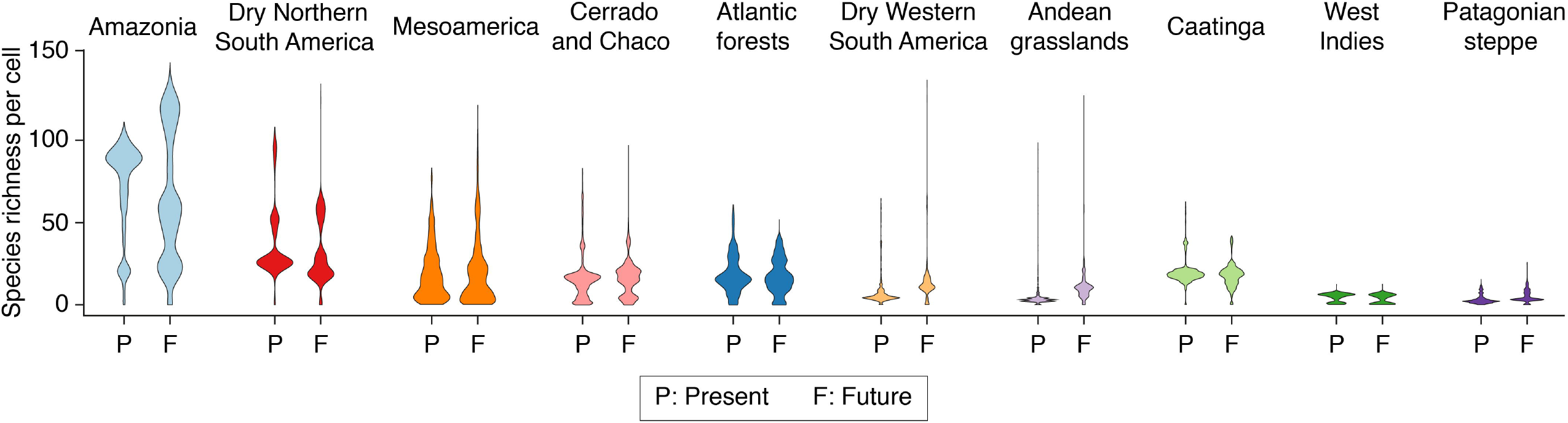
Current (Present) and projected (future) richness of bat species per cell in the Neotropical regions under the business-as-usual (B.A.U) scenario of climate and land-use changes and assuming limited dispersal (bat species can disperse only across analog environment).

**Table 2:** Potential change in species richness of Neotropical bats for ecoregions at the local scale. Numbers represent the mode of species richness for all cells in each ecoregion in the present and projected under the business-as-usual (B.A.U) scenario of climate and land-use changes and assuming limited dispersal (bat species have the ability to disperse only over analog environment).

| Ecoregions | Present | Future |  |
| --- | --- | --- | --- |
|  | Cell<br>richness | Cell<br>richness | Gain (+) Loss (-) |
| **<u>Forested Wet</u>** |  |  |  |
| Amazonia | 85 | 59 | -26 |
| Atlantic Forests | 16 | 17 | 1 |
| **<u>Open Dry</u>** |  |  |  |
| Andean Grasslands | 3 | 10 | +7 |
| Caatinga | 18 | 18 |  |
| Cerrado and Chaco | 15 | 17 | +2 |
| Dry Northern South<br>America | 27 | 26 | -1 |
| Dry Western South<br>America | 4 | 11 | +7 |
| Patagonian Steppe | 2 | 4 | +2 |
| **<u>Mixed</u>** |  |  |  |
| Mesoamerica | 15 | 16 | +1 |
| West Indies | 6 | 5 | -1 |

**FIG. 4.**
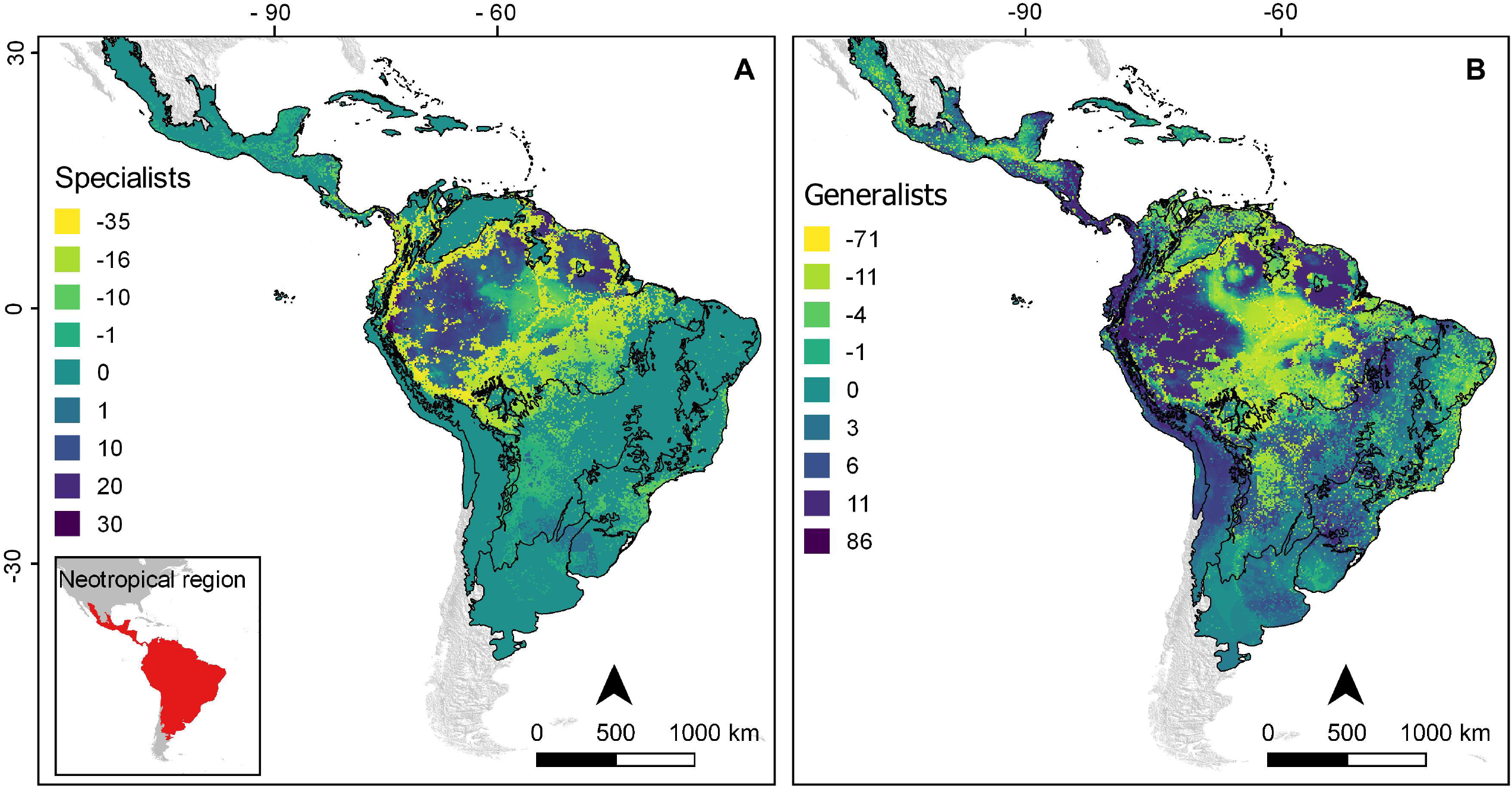
Projected change in richness of (A) environment-specialist and (B) environment-generalist bat species under the business-as-usual scenario of climate and land use changes (B.A.U). Dispersal is assumed to be limited so that bat species have the ability to disperse only across analog environment.

Our projections indicate different trophic guilds will be impacted differently by climate and land use change (Fig. 5). Within most ecoregions, the number of frugivorous bats projected to undergo range reductions is more than twice as large as the number of species projected to undergo expansions (Appendix S4-S13). For other guilds, patterns are less consistent. For instance, in the Caatinga (Appendix S6) and Cerrado/Chaco (Appendix S8) regions more insectivores are projected to undergo range reduction than expansion, while in the Dry Western (Appendix S10) and Patagonian steppe (Appendix S13) most insectivores are projected to expand in range. Two sanguivorous bats (*Diaemus youngi* and *Diphylla ecaudata*) are projected to expand their ranges into Dry Western and Patagonian steppe respectively (Appendix S10 and S13) and five Neotropical endemic bat species *Phyllostomus latifolius* (omnivore, Dry Northern South America), *Dryadonycteris capixaba* and *Lonchophylla mordax* (nectarivore, Atlantic Forest), *Chilonatalus micropus* (insectivore, West Indies) and *Eptesicus innoxius* (insectivore, Dry Western South America) are predicted to become globally extinct by the combined effects of climate and land use changes (Table 1).

**FIG. 5.**
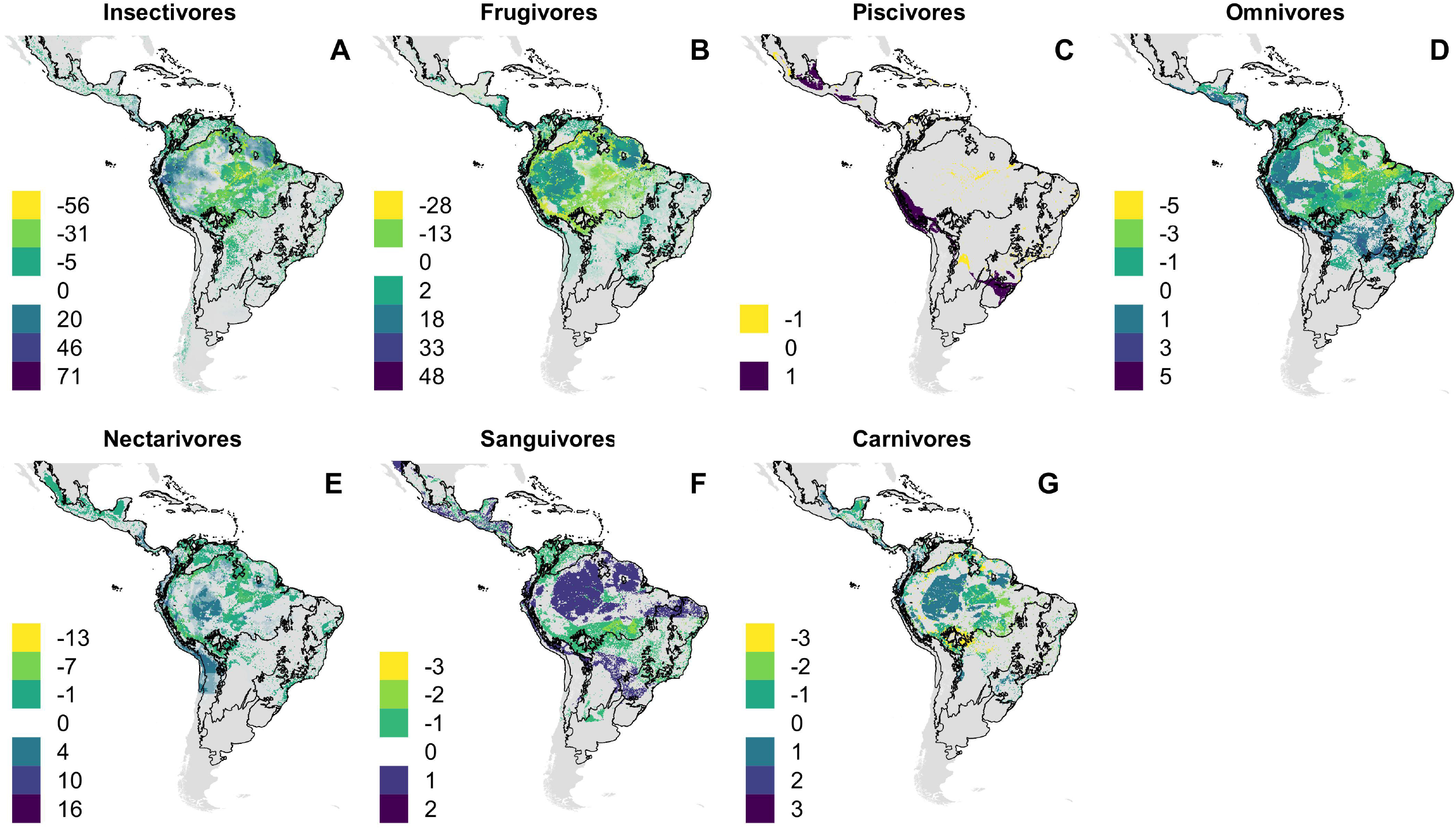
Projected change in richness of bat species according to feeding habits and under the business-as-usual scenario of climate and land use changes (B.A.U). Dispersal is assumed to be constrained so that bat species have ability to disperse only over land cover analog to current occurrence patterns. (A) Insectivores, (B) Frugivores, (C) Piscivores, (D) Omnivores, (E) Nectarivore, (F) Sanguivores and (G) Carnivores.

## Discussion

### The magnitude of range shift

This is the first effort to evaluate the possible future consequences of two of the most important drivers of current global environmental change - land use and climate change - on Neotropical bats. Focusing on the more realistic scenario of socio-economic development (B.A.U), our projections suggest substantial future declines in environmental suitability for the Neotropical bat fauna, especially for bat species that have specialised on a specific type of environment. The widespread loss of specialised species in our projections was compensated by the expansion of generalised species that forage in open areas and commonly establish large colonies in manmade structures (Gonçalves et al., 2017). The balance between the loss of habitat specialists and expansion of generalists may lead to little net change of species richness at the local level, but greater homogeneity in species composition. Functional homogenisation is considered a global prominent forms of biodiversity impoverishment induced by recent environmental change (Devictor et al., 2008; Clavel et al., 2011). Species that exhibit morphological and functional specialization on specific conditions are more sensitive to changes because unique traits restrict them to only a narrow set of environments and resources (Clavel et al., 2011; Newbold et al., 2018; Staude et al., 2020). In addition, the use of a restricted range of habitats implies that habitat-specialist bat species are unable or unlikely to cross-gaps of unsuitable environment to colonize isolated suitable patches. Therefore, habitat specialists are not only threatened by changes in their current habitat, but also by the inability to access novel suitable habitat.

### Neotropical ecoregions with higher projected impacts

Several Neotropical bat species may be forced to shift their range to refuges that are less vulnerable climate change, venturing into regions of different elevation and/or latitudes while facing new environmental challenges in response to the land use and climate change. Our projections showed that areas nearby the Andean mountains (Northern Andean) and the Guiana Highlands would be considered climate change refugia for several Neotropical bat species (Fig. 4; Fig. 5). Yet, Climate-driven migrations will only allow occupancy of newly suitable climates if permeable migratory routes allow dispersal across landscapes (Engler et al., 2012). When we considered dispersal barriers, our analyses revealed no net gains of species in Amazonia although this region is projected to harbour suitable conditions for many species. Reduced accessibility due to the ‘Arch of Deforestation’ (Sales et al., 2019), a region of high agricultural pressure from cattle-ranching and soy plantations, restricts climate-driven movements of forest specialist species from the regions south of the Amazon basin and vice-versa.

Our models predicted higher gains of species in the Dry Western South America (61 species, increasing potential richness in 60%) located in the Andes foothills at mid-elevation (1000-2500m). This means that there is a great potential for Neotropical bat species immigration at these elevations. The major factors driving this trend are the upslope dispersal of lowland species in response to global changes combined with projections of low changes in land use in the region. Such upslope movements are predicted by the hypothesis of upward and upslope shifts in species ranges due to elevated temperatures (Colwell et al., 2008) and have been reported for birds (Bender et al., 2019) and dung beetles (Larsen, 2012) in the same region and for bats in the Mountains of northern Costa Rica (LaVal 2004, Arias-Aguilar et al., 2020).. Our findings thus highlight the potential of mountains and highlands to act as refugia for bats occurring in low elevation over time.

Conversely, our models predicted high losses of species from the regional species pool in the Andean Grassland (22 species, representing 13% of total richness) located at high-elevation (3200-3500m). Yet, all species currently occurring in this region were projected to persist at the continental scale, as long as they can disperse to suitable environment (Table 1). This could be a consequence of the wide elevational ranges of species at such altitude which is consistent with Rapoport’s rule that species at high elevations tend to have wider elevational ranges, covering several elevational levels, whereas species in the lowlands tend to have narrow elevational ranges (Stevens, 1992). The greatest proportional loss in species richness was found for the West Indies, 27% of the species (7 out of the 26) and the Caatinga, 22% (17 out of 79 species). Species in these regions are projected to be subjected to non-analogue climates and habitat loss, which combined with dispersal limitation, will prevent occupancy of neighbouring suitable environment. These species are thus expected to undergo severe range contractions because of their inability to colonize the novel suitable environments that may surround their current ranges. Our projections also showed high species turnover in Cerrado and Chaco (Table 1), where losses may be partially compensated by gains.

Five endemic bat species that occupy small areas in the Neotropics are predicted to become globally extinct (Table 1), but this number may be higher since our conservative analysis is restricted to species whose range size was larger than 30 cells (Appendix S1). Neotropical endemic bat species and/or species that occupy such a small area, may have entire populations exposed to physiological stress or resource shortage even under small environmental variation (Newbold et al., 2018; Staude et al., 2020). Those narrow-ranged species are also vulnerable to non-climatic stressors, such as population stochasticity, low variability and Allee effect, that impact small populations (Newbold et al., 2018).

### Shifts in ecological functions

Our analyses revealed that combined effects of land use and climate change are likely to favour many animalivorous bat species that forage in unobstructed areas outside or above forested sites and easily establish high-densities colonies in manmade structures (Gonçalves et. al. 2017; Farneda et al., 2019). Those are typical attributes of molossid and some phyllostomid bats (Simmons, 2005), which may allow species to increase their ranges by colonizing other ecoregions in response to changes. Bat species with these characteristics are usually evaluated with a lower risk of extinction (IUCN, 2020), which is consistent with their success to persist within modified landscapes and under variable climatic conditions. However, several phytophagous bats (nectarivores and frugivores) are projected to experience major reductions in their ranges. Species that rely on the vegetation as food sources and natural roosting places are more likely to be impacted by land use change (Popa-Lisseanu and Voigt, 2009; Moussy et al., 2013; Sherwin et al., 2013; Gonçalves et al., 2017; Farneda et al., 2019). The interplay between species dispersal abilities and its ecological requirements will shape the species-specific responses to environmental change.

These uneven outcomes for different trophic guilds may impact ecosystem functions locally. As species expand or retract, their interactions are also redistributed. The replacement of phytophagous bats by insectivore species may result in local declines in pollination and seed dispersal of many plants. Likewise, range reduction of carnivorous and piscivorous bats forecasted for some regions may impair the control of prey populations and transport of nutrients across the landscape (Gonçalves et al., 2017). Range expansions of a species may also result in the expansion of its pathogens, including those associated with zoonotic diseases (Gonçalves et al., 2020). The projected range expansion of two sanguivorous bats into Dry Western and Patagonian steppe for instance, could increase the probability of transmission of rabies and other saliva-borne viruses (Olival et al., 2017).

### Implications for species extinction risk and conservation biogeography

Based on the expected changes in climatic conditions in the Neotropics, we argue that the most effective way to protect these species will be by increasing landscape connectivity through habitat restoration, connecting native vegetation fragments and forest patches with broad corridors (Arroyo-Rodríguez et al., 2020). Despite the potential of bats for large-scale species migration, reserves should also remain as a component of conservation strategy under land use and climate changes. Creating new protected areas, especially surrounding climate change refugia, and preserving existing ones will also act to protect high quality habitat, reducing extinction risk as well as providing suitable regions for colonizing species or stepping stones for species on the move. A mosaic of land uses and less damaging landscape practices, such as low intensity forestry, may also provide some opportunity for persistence or migration (Arroyo-Rodríguez et al., 2020). Enhancing existing linear habitat features, such as river banks, hedgerows and embankments may improve connectedness without actively creating new corridors (Sales et al., 2019).

Proactive conservation can be applied in all ecoregions but should especially focus in the Northern Andean mountains, Guiana highlands and Mesoamerica mountains as these regions still retain a high percentage of natural habitat where the biota is relatively intact (Antonelli et al., 2018; Ribeiro et al., 2017; Sales et al., 2019; Fragoso et al., 2019). In these regions, large-scale conservation, such as the protection of a large extension of land, might be achieved with relatively low investment. Due to the highest loses of bat species projected for the West Indies (specially the Caribbean dry forest), Andean grassland, and for the Caatinga and Cerrado/Chaco regions we stress the need for efficient measures to halt habitat loss and fragmentation and continued monitoring of species responses to climate change.

Fine-scale reactive conservation should be applied for long-term monitoring of climate change responses by five endemic bat species that are predicted to become extinct and also species that occupy small areas and in the Neotropics. Monitoring would also be important to help detecting the functional shifts related to expansion/retraction of bats from different guilds projected here, and is especially important considering the projected range expansion of two vampire bats into new ecoregions (Gonçalves et al., 2021), which might be of public health concern (Olival et al., 2017). We caution that our projections considering multiple species simultaneously have been designed for large scale assessment of broad distribution patterns and did not account for species-specific physiological or behavioural responses to small scale environmental and landscape properties. Thus, conservation planning focused at individual species should not rely exclusively on such models, which can be used as a starting point, but should be combined with fine-grained field data as well as a more in-depth analysis of how different climate variables and biotic contexts affect fitness (Mawdsley et al., 2009).

## Conclusions

Overall, our results provide new insights to guide landscape management, policy and practice to maintain or enhance bat diversity and their ecological functions in the Neotropics. A main highlight of our projections is that, despite the low number of habitat specialist species observed in land use/climate change-influenced area, increasing landscape connectivity and maintenance of old-growth forests is crucial for Neotropical bat ecosystem services conservation in the future. Long-term monitoring programs will be important to track the combined impacts of climate and land use changes on Neotropical bats and provide conservationist with information to evaluate the effectiveness of conservation plans (Sherwin et al., 2013). Furthermore, better determining the dispersal abilities, realized niche width and physiological tolerances at the species level will be particularly important for understanding the vulnerability of Neotropical bats to climate and land use change and their ability to persist or redistribute in response to the changing environment (Mawdsley et al., 2009; Zamora-Gutierrez et al., 2018).

## Supplementary Material

Additional information (Appendix S1-S13) and R code are available online. The authors are solely responsible for the content and functionality of these materials. Queries (other than absence of the material) should be directed to the corresponding author. Supplementary Information is also available on GitHub (github.com/fhmgoncalves).

## Acknowledgements

FG, MG and MMP were supported by (FAPESP) São Paulo Research Foundation (Grant 2017/24252-0, 2019/00648-7, 2019/25478-7). LS was funded by the PNPD (Programa Nacional de Pós-Doutorado, in Portuguese) at the Campinas State University (UNICAMP). This study was financed in part by the Coordenação de Aperfeiçoamento de Pessoal de Nível Superior - Brasil (CAPES) - Finance Code 001.

