## Appendix S1 for "Combined impacts of climate and land use change and the future restructuring of Neotropical bat biodiversity"

**Methods**

**Distribution information across the Neotropics**

The Neotropics harbor the highest diversity of bats in the world: nine families (Furipteridae, Mormoopidae, Noctilionidae, Phyllostomidae, Thyropteridae, Emballonuridae, Vespertilionidae, Molossidae and Natalidae) comprising 83 genera and around 280 species that can be assigned to seven different trophic guilds (carnivore, insectivore, frugivore, nectarivore, omnivore, piscivore and sanguivore) (Mickleburgh et al. 2002). To model the potential effects of climate and land use changes on bat diversity and avoid overestimation, we modelled only Neotropical bat species whose range size was larger than 30 cells totalling 228 bat species (ca. 80% of all species recorded in the Neotropics) distributed across the nine bat families: Phyllostomidae (138); Molossidae (31); Vespertilionidae (32); Emballonuridae (16); Mormoopidae (5); Natalidae (5); Thyropteridae (2); Noctilionidae (2) and Furipteridae (1). We focused our analyses on the Neotropics ranging from northern Mexico to central Argentina and including the whole Caribbean (Morrone 2014). There is no general agreement on how Neotropical ecoregions should be classified, with sometimes substantial disparity between specialist based and data-driven strategy (Edler et al. 2017), as well as variation in the number and definition of boundaries of ecoregions based on the same data but derived from different methods (Vilhena and Antonelli, 2015). Taking that into account, we adopted in our analyses the Neotropical ecoregions proposed by Antonelli et al. (2018) as both studies have focused on dispersal processes. Antonelli et al. (2018) used the classification of terrestrial biomes and ecoregions by Olson et al. (2001), which is widely used in ecology and biogeography, but reduced the number of biomes in order to highlight the major ecosystems that are well defined according to geological history and/or geographical barriers and to reduce the risk of erroneous geographic distribution of species into each ecoregion (see Antonelli et al. (2018) for more details on how they splited geographically disjunct biomes).

The available information on the occurrence of bats is highly biased with wide spatial gaps in regions such as the Amazon and southern South America. Therefore instead of relying upon occurrence points we used the range maps from the IUCN database (www.iucnredlist.org, date of access: August 08th, 2020) as the basis for our distribution modelling. To avoid model overfitting and reduce commission errors, instead of considering sampling points within the full species range, we randomly sampled points within each species’ distribution according to range size, following methods recently described (Sales et al. 2019, 2020). To do so, we first rasterized the IUCN range maps into a gridded file of 10’ resolution (approximately 0.17 degree of lat/long) and, to be conservative and avoid overestimation, we restricted our analysis to species whose range size was larger than 30 cells. For species whose range size was larger than 1000 cells, we sampled 12.5% of these cells, thus considered “presences” in the rasterized map. For species with range size varying from 501 to 1000 and 101 to 500 we sampled 25% and 50% of the cells within species ranges. Finally, for species with less than 100 cells within the total range we used all cells as occurrences to calibrate distribution models.

**Climate Data**

Climate information was obtained as 10’ resolution gridded raster files for land areas, derived from interpolation of worldwide ground weather stations, available in CliMond (https://www.climond.org/). We downloaded climate data as and encompassing 30 years of measurements, from 1961 – 1990. Input data for the CliMond baseline climatology was obtained from WorldClim (Hijmans et al. 2005), CRU CL1.0 and CRU CL2.0 (New et al. 2002). We chose CliMond as a global climate repository because it includes humidity data and 35 additional bioclimatic variables that are climatic relevant aspects that influence, direct or indirectly, bat species distribution and survival in the Neotropics (Sherwin et al. 2013), as well as a wider range of timesteps for future forecasts. These intermediate timesteps on climate forecasts allow incorporation of climatic bridges between present and future climate analogs, with the explicit incorporation of species’ dispersal abilities (Sales et al. 2019, 2020).

We project future climates based on two Atmospheric Ocean Global Circulation Models (AOGCMs): CSIRO (Gordon et al. 2002) and MIROC-H (Chikamoto et al. 2013), which are widely used in climate change literature and known to produce reliable information. They build upon the expected trajectories of emission of greenhouse gases and sulphate aerosol from the last assessment cycle of the International Panel on Climate Change (IPCC-AR5) (IPCC 2014). The modeled scenarios depend on global human demographic, economic and technological development so that the Mitigation, A1B scenario, estimates a temperature increase of 1.7 – 4.4ºC by the end of the 21^st^ century, whereas the, Business-as-usual, A2 scenario, forecasts a 2.0 – 5.5ºC temperature increase. Thus, these roughly correspond to the representative concentration pathways *RCP6.0* and *RCP8.5* from the IPCC-AR5 (IPCC 2014). Those forecasts indicate temperature increases above the 1.5ºC threshold recommended to avoid the deleterious effects of climate change on ecosystems and human well-being (IPCC 2019) but are congruent with observed changes from years 2000-2010 (IPCC 2019).

Among the 35 CliMond predictors that reflect plausible constraints on energy, water and temperature, all of which can contribute to determine bat distributions in the Neotropics, we selected only those not subjected to high multicollinearity (Graham 2003) because the use of an excessive number of predictors often leads to over-fitted models, with limited predictive ability outside the training domain (Kriticos et al. 2012). Although other factors might also affect species distribution, several studies indicate that climate is the main driver of species distribution at large spatial scales (Ribeiro et al. 2016).

**Land-use and land-cover**

To obtain projections of changes in land use we used a global land-use land-cover (LULC) model based on human-environment interactions (Li et al. 2017). This is one of the only models that provide global projections of LULC change for scenarios of future human development at a very high resolution (1km², which was later aggregated into 18km² with a resampling procedure with “nearest neighbor” interpolation, function *resample()* from the *raster* R package – Hijmans 2012), incorporating human-environment dynamics as the main driver of landscape changes. The model consists of one “top-down” and one “bottom-up” component. The first refers to an economic modeling using a downscaled version of the Integrated Model to Assess the Global Environment (IMAGE), an agro-economic method of estimating land-use allocation, considering a range of factors such as climate, soil and population density. In IMAGE, LULC changes until the required regional production of crops and grass is met, according to different scenarios of human development (IPCC, 2019), and defines dispersal constraints of the next component. The “bottom-up” element of the model includes a future land use simulation (the FLUS model - Liu et al., 2017) based on a cellular automata model. The occurrence probability surfaces of each land-cover type were determined by biophysical and socioeconomic drivers, where self-adaptive inertia, competition and interactions among land-use types defined LULC changes (Li et al. 2017).

We used LULC projections referred to scenarios A1B and A2, the same ones we chose for obtaining climate data at CliMond. The combination of the land-cover classes associated to a certain species habitat was, therefore, considered an environmental filter and applied as a mask onto climate-based predictions of potential distribution. The resulting LULC map was composed of six classes (forest, savanna, grassland, farmland, urban, and barren) for year 2010 and projections for 2050 and 2100. By doing so, we are explicitly considering species-specific habitat sorting mechanisms which prevent occupancy of non-analog land cover. These masks were, then, applied to all timesteps of climate-based predictions of distribution potential. In the absence of decadal LULC predictions, we combined the closest possible information for climate and land-cover.

**Modelling occupancy dynamics and dispersal**

We simulate dispersal using the cellular automata model implemented in the R package MigClim (Engler et al., 2012). The basic unit of this model is a cell that is occupied or not (here considered those cells that contain suitable climate and land cover) at time *t*_initial_, defining the initial distribution. Occupancy dynamics are, then, driven by environmental changes and species dispersal and colonization ability (Engler et al. 2012; Engler and Guisan, 2009). If environmental conditions remain suitable across consecutive time steps (*t*_initial_, *t* +1, *t* + 2, …, *t*_final_), occupied cells suffer no changes. In case the cell becomes unsuitable (given climate or land-cover changes), occupied cells become empty (decolonized). An unsuitable cell in time *t* (target) can be colonized if a) environmental conditions became suitable in time *t+1*, and b) the distance from the closest source cell (*d*) is smaller than the species maximum dispersal ability (Engler and Guisan, 2009), here considered one cell per year, due to the absence of species-specific information.

We then included the effect of land-use changes, by including non-analog land cover cells as barriers to dispersal. Barrier cells are considered permanently unsuitable for species, thus colonization is not possible, but they also prevent dispersal through them (Engler et al. 2012). We modulated the strength of non-analog climates as barrier (strong *vs* weak) to dispersal according to a general response of habitat specialist *vs* generalist bat species to land use change and fragmentation (Büchi and Vuilleumier, 2014). Habitat specialist species were all those species whose occurrence was restricted to a single major land cover type (e.g. forest), regardless the specific category of land cover (88 species). Accordingly, we consider a habitat generalist any species whose occurrence is not restricted to a single major land cover type (e.g. forest and farmland) (140 species). Non-analog land cover was then included as strong barrier for habitat specialist bats but a weak barrier to habitat generalist bats. The difference in barrier strength lies on the dispersal between two diagonally adjacent barrier cells, allowed if the barrier is weak but not possible for strong barriers (Engler et al. 2012). Range shift was calculated as the percent variation in the number of suitable cells for a given species, comparing the current and future potential distribution (Range*_future_* − Range*_current_*) × 100/Range*_current_*.

The incorporation of dispersal constraints results in a map of dispersal-restricted potential distribution, where occupancy dynamics are partitioned in space. Cells predicted to be suitable and occupied in all timesteps are regarded as climate refugia. Those newly suitable target cells that are accessible via migration from source cells indicate zones of potential migration. Cells that become suitable but are out of reach indicate dispersal limitation regions. Finally, cells that are suitable in the present but will become unsuitable in the future are considered cells with non-analog environment, which are likely to expose populations to conditions for which species are not adapted to (Ribeiro et al. 2016).

We used the following parameterization of function “MigClim.migrate” for all dispersal simulations. The function argument hsMap included the habitat suitability models for intermediate timesteps (years 2030, 2050 and 2070); envChgSteps and dispSteps, i.e. the number of environmental change steps and the number of dispersal steps were set to three (the number of intermediate timesteps). The argument barrier included a raster file with the land-use and land-cover types listed as unsuitable for each species, and barrierType was defined as “weak” or “strong”, depending on habitat generalization or specialization, respectively. Finally, replicateNB was set to three full simulations of dispersal per species. For the remaining arguments we used the function default.

**Appendix S2.** Information about each studied bat species regarding habitat use, feeding habits and distribution models. AUC is the area under the receiver operating characteristic curve computed for the species distribution models. Current and future potential ranges refer to the number of suitable cells in the present and projected to be occupied in the future under the limited dispersal scenario.

| **Species name** | **Habitat use** | **Feeding habit** | **Model**  **AUC** | **Current potential range** | **Future (A1B) potential range** | **Future (A2) potential range** |
| --- | --- | --- | --- | --- | --- | --- |
| *Ametrida centurio* | specialist | frugivore | 0.75 | 10389 | 9487 | 4080 |
| *Anoura aequatoris* | specialist | nectarivore | 0.85 | 967 | 2387 | 1283 |
| *Anoura caudifer* | specialist | nectarivore | 0.69 | 8589 | 15717 | 8171 |
| *Anoura cultrata* | generalist | nectarivore | 0.83 | 1451 | 4380 | 4180 |
| *Anoura fistulata* | specialist | nectarivore | 0.91 | 183 | 1279 | 299 |
| *Anoura geoffroyi* | generalist | nectarivore | 0.68 | 11629 | 21316 | 18524 |
| *Anoura latidens* | generalist | nectarivore | 0.86 | 1464 | 4257 | 3983 |
| *Anoura luismanueli* | generalist | nectarivore | 0.94 | 111 | 375 | 29 |
| *Anoura peruana* | generalist | nectarivore | 0.82 | 2598 | 5304 | 5265 |
| *Artibeus aequatorialis* | specialist | frugivore | 0.87 | 273 | 805 | 191 |
| *Artibeus amplus* | specialist | frugivore | 0.65 | 760 | 4508 | 2339 |
| *Artibeus concolor* | specialist | frugivore | 0.71 | 11868 | 12315 | 6712 |
| *Artibeus fimbriatus* | specialist | frugivore | 0.8 | 745 | 324 | 7 |
| *Artibeus fraterculus* | generalist | frugivore | 0.86 | 177 | 107 | 66 |
| *Artibeus jamaicensis* | generalist | frugivore | 0.85 | 3061 | 3237 | 3359 |
| *Artibeus lituratus* | generalist | frugivore | 0.69 | 24769 | 23680 | 21236 |
| *Artibeus obscurus* | generalist | frugivore | 0.64 | 28602 | 27024 | 19368 |
| *Artibeus planirostris* | generalist | frugivore | 0.68 | 14491 | 18031 | 16458 |
| *Carollia benkeithi* | specialist | frugivore | 0.79 | 3639 | 2351 | 89 |
| *Carollia brevicauda* | generalist | frugivore | 0.68 | 26304 | 24649 | 22798 |
| *Carollia castanea* | generalist | frugivore | 0.68 | 4762 | 12198 | 10827 |
| *Carollia manu* | specialist | frugivore | 0.85 | 12 | 81 | 1 |
| *Carollia perspicillata* | generalist | frugivore | 0.66 | 20034 | 17965 | 8892 |
| *Centronycteris centralis* | specialist | insectivore | 0.71 | 4279 | 12454 | 7594 |
| *Centronycteris maximiliani* | specialist | insectivore | 0.73 | 9723 | 12955 | 7934 |
| *Centurio senex* | generalist | frugivore | 0.78 | 3461 | 6837 | 6474 |
| *Chilonatalus micropus* | specialist | insectivore | 0.88 | 1 | 0 | 0 |
| *Chiroderma doriae* | generalist | frugivore | 0.8 | 1385 | 1982 | 1960 |
| *Chiroderma salvini* | generalist | frugivore | 0.71 | 7351 | 11499 | 6815 |
| *Chiroderma trinitatum* | generalist | frugivore | 0.71 | 16531 | 13822 | 6483 |
| *Chiroderma villosum* | specialist | frugivore | 0.72 | 18081 | 16261 | 8072 |
| *Choeroniscus godmani* | generalist | frugivore | 0.74 | 4498 | 7565 | 6306 |
| *Choeroniscus minor* | generalist | frugivore | 0.75 | 13309 | 21310 | 20427 |
| *Choeroniscus periosus* | specialist | nectarivore | 0.96 | 156 | 315 | 60 |
| *Chrotopterus auritus* | generalist | carnivore | 0.65 | 22817 | 23730 | 21360 |
| *Cormura brevirostris* | specialist | insectivore | 0.72 | 16176 | 12763 | 5886 |
| *Cynomops abrasus* | specialist | insectivore | 0.68 | 16753 | 14684 | 7263 |
| *Cynomops greenhalli* | generalist | insectivore | 0.76 | 8677 | 11281 | 9496 |
| *Cynomops milleri* | specialist | insectivore | 0.77 | 4753 | 6438 | 3067 |
| *Cynomops paranus* | specialist | insectivore | 0.75 | 12749 | 15606 | 8772 |
| *Cynomops planirostris* | specialist | insectivore | 0.65 | 15990 | 16305 | 8495 |
| *Cyttarops alecto* | generalist | insectivore | 0.89 | 1509 | 3671 | 3223 |
| *Dermanura anderseni* | generalist | frugivore | 0.75 | 9092 | 10177 | 8408 |
| *Dermanura bogotensis* | specialist | frugivore | 0.71 | 488 | 6191 | 3826 |
| *Dermanura cinerea* | generalist | frugivore | 0.74 | 10576 | 14372 | 14068 |
| *Dermanura glauca* | generalist | frugivore | 0.71 | 5663 | 18767 | 16409 |
| *Dermanura gnoma* | generalist | frugivore | 0.66 | 17575 | 18637 | 15055 |
| *Dermanura phaeotis* | generalist | frugivore | 0.69 | 7314 | 16348 | 14720 |
| *Dermanura rava* | specialist | frugivore | 0.83 | 415 | 411 | 78 |
| *Dermanura rosenbergi* | specialist | frugivore | 0.92 | 219 | 235 | 46 |
| *Dermanura tolteca* | generalist | frugivore | 0.86 | 1401 | 1418 | 1652 |
| *Dermanura watsoni* | generalist | frugivore | 0.79 | 991 | 1479 | 756 |
| *Desmodus rotundus* | generalist | sanguivore | 0.65 | 9797 | 7819 | 8794 |
| *Diaemus youngi* | generalist | sanguivore | 0.68 | 21589 | 22432 | 19764 |
| *Diclidurus albus* | generalist | insectivore | 0.69 | 21152 | 20976 | 18172 |
| *Diclidurus ingens* | generalist | insectivore | 0.73 | 3692 | 1926 | 1513 |
| *Diclidurus isabella* | generalist | insectivore | 0.84 | 2479 | 3335 | 2663 |
| *Diclidurus scutatus* | generalist | insectivore | 0.86 | 3557 | 10157 | 9479 |
| *Diphylla ecaudata* | generalist | sanguivore | 0.69 | 19187 | 31776 | 23235 |
| *Dryadonycteris capixaba* | specialist | nectarivore | 0.92 | 1 | 0 | 0 |
| *Enchisthenes hartii* | specialist | frugivore | 0.72 | 3585 | 11700 | 5424 |
| *Eptesicus brasiliensis* | specialist | insectivore | 0.68 | 19105 | 17788 | 8995 |
| *Eptesicus chiriquinus* | specialist | insectivore | 0.77 | 372 | 11127 | 5859 |
| *Eptesicus diminutus* | generalist | insectivore | 0.8 | 2718 | 5593 | 4298 |
| *Eptesicus furinalis* | generalist | insectivore | 0.62 | 26590 | 26093 | 23207 |
| *Eptesicus fuscus* | generalist | insectivore | 0.75 | 27605 | 23917 | 24247 |
| *Eptesicus innoxius* | specialist | insectivore | 0.87 | 45 | 8 | 0 |
| *Eptesicus taddeii* | specialist | insectivore | 0.85 | 73 | 158 | 4 |
| *Eumops auripendulus* | generalist | insectivore | 0.65 | 39769 | 41445 | 40704 |
| *Eumops bonariensis* | generalist | insectivore | 0.87 | 1357 | 1989 | 1749 |
| *Eumops dabbenei* | generalist | insectivore | 0.71 | 6030 | 18814 | 18591 |
| *Eumops delticus* | specialist | insectivore | 0.77 | 2073 | 5315 | 3271 |
| *Eumops glaucinus* | generalist | insectivore | 0.69 | 15524 | 18078 | 15786 |
| *Eumops hansae* | generalist | insectivore | 0.75 | 16944 | 17973 | 11024 |
| *Eumops maurus* | generalist | insectivore | 0.77 | 3208 | 11688 | 10656 |
| *Eumops nanus* | generalist | insectivore | 0.76 | 62 | 566 | 535 |
| *Eumops patagonicus* | generalist | insectivore | 0.86 | 2751 | 3504 | 2750 |
| *Eumops perotis* | generalist | insectivore | 0.73 | 18056 | 23324 | 21109 |
| *Eumops trumbulli* | generalist | insectivore | 0.77 | 11347 | 10214 | 8820 |
| *Furipterus horrens* | generalist | insectivore | 0.69 | 14484 | 13841 | 6929 |
| *Gardnerycteris crenulatum* | generalist | insectivore | 0.69 | 29703 | 31346 | 29264 |
| *Glossophaga commissarisi* | specialist | nectarivore | 0.77 | 7119 | 12054 | 6741 |
| *Glossophaga longirostris* | generalist | nectarivore | 0.67 | 2988 | 6827 | 4070 |
| *Glossophaga soricina* | generalist | nectarivore | 0.72 | 40449 | 40780 | 40090 |
| *Glyphonycteris behnii* | generalist | insectivore | 0.84 | 2725 | 7264 | 5073 |
| *Glyphonycteris daviesi* | specialist | insectivore | 0.8 | 10385 | 8269 | 4883 |
| *Glyphonycteris sylvestris* | generalist | insectivore | 0.81 | 12442 | 16007 | 8909 |
| *Histiotus humboldti* | specialist | insectivore | 0.89 | 250 | 1509 | 579 |
| *Histiotus laephotis* | specialist | insectivore | 0.86 | 746 | 864 | 59 |
| *Histiotus macrotus* | generalist | insectivore | 0.81 | 874 | 3823 | 3825 |
| *Histiotus magellanicus* | specialist | insectivore | 0.88 | 570 | 629 | 171 |
| *Histiotus montanus* | generalist | insectivore | 0.79 | 7952 | 11152 | 10760 |
| *Histiotus velatus* | generalist | insectivore | 0.78 | 2166 | 3198 | 2604 |
| *Lampronycteris brachyotis* | generalist | omnivore | 0.79 | 9173 | 13920 | 12736 |
| *Lasiurus blossevillii* | generalist | insectivore | 0.72 | 43199 | 54697 | 53622 |
| *Lasiurus cinereus* | generalist | insectivore | 0.69 | 44687 | 49405 | 49562 |
| *Lasiurus ega* | generalist | insectivore | 0.67 | 20821 | 23824 | 21047 |
| *Lasiurus egregius* | specialist | insectivore | 0.85 | 2071 | 8851 | 5816 |
| *Lasiurus salinae* | specialist | insectivore | 0.87 | 415 | 830 | 750 |
| *Lasiurus varius* | generalist | insectivore | 0.88 | 947 | 934 | 518 |
| *Leptonycteris curasoae* | generalist | nectarivore | 0.78 | 439 | 439 | 57 |
| *Lichonycteris degener* | specialist | nectarivore | 0.77 | 6133 | 15514 | 8802 |
| *Lichonycteris obscura* | generalist | nectarivore | 0.8 | 1371 | 2104 | 2197 |
| *Lionycteris spurrelli* | generalist | nectarivore | 0.74 | 19907 | 21219 | 19871 |
| *Lonchophylla concava* | generalist | nectarivore | 0.85 | 427 | 1310 | 1432 |
| *Lonchophylla handleyi* | generalist | nectarivore | 0.86 | 218 | 878 | 701 |
| *Lonchophylla mordax* | generalist | nectarivore | 0.95 | 1 | 0 | 0 |
| *Lonchophylla orienticollina* | specialist | nectarivore | 0.73 | 143 | 124 | 2 |
| *Lonchophylla robusta* | generalist | nectarivore | 0.75 | 1430 | 1635 | 1585 |
| *Lonchophylla thomasi* | generalist | nectarivore | 0.72 | 18318 | 14443 | 11727 |
| *Lonchorhina aurita* | generalist | insectivore | 0.69 | 20343 | 21341 | 19001 |
| *Lonchorhina fernandezi* | generalist | insectivore | 0.98 | 17 | 0 | 3 |
| *Lonchorhina inusitata* | generalist | insectivore | 0.78 | 6982 | 4408 | 2456 |
| *Lonchorhina marinkellei* | generalist | insectivore | 0.92 | 14 | 0 | 4 |
| *Lonchorhina orinocensis* | generalist | insectivore | 0.82 | 951 | 5490 | 3185 |
| *Lophostoma brasiliense* | generalist | insectivore | 0.73 | 20840 | 26384 | 26285 |
| *Lophostoma carrikeri* | generalist | insectivore | 0.78 | 8130 | 10916 | 7327 |
| *Lophostoma silvicolum* | specialist | insectivore | 0.75 | 13508 | 12579 | 6933 |
| *Macrophyllum macrophyllum* | specialist | insectivore | 0.65 | 19112 | 17670 | 8673 |
| *Mesophylla macconnelli* | specialist | frugivore | 0.73 | 17336 | 13418 | 6013 |
| *Micronycteris brosseti* | specialist | insectivore | 0.82 | 1316 | 3569 | 2065 |
| *Micronycteris hirsuta* | generalist | insectivore | 0.79 | 14554 | 15913 | 13159 |
| *Micronycteris megalotis* | generalist | insectivore | 0.63 | 20149 | 17797 | 14367 |
| *Micronycteris microtis* | generalist | insectivore | 0.77 | 9640 | 16433 | 14838 |
| *Micronycteris minuta* | generalist | insectivore | 0.66 | 20912 | 21379 | 18873 |
| *Micronycteris schmidtorum* | specialist | insectivore | 0.78 | 11457 | 13414 | 8215 |
| *Mimon bennettii* | generalist | insectivore | 0.64 | 13405 | 23884 | 17153 |
| *Mimon cozumelae* | generalist | insectivore | 0.71 | 898 | 6663 | 4957 |
| *Molossops mattogrossensis* | generalist | insectivore | 0.72 | 9613 | 13817 | 7360 |
| *Molossops temminckii* | generalist | insectivore | 0.62 | 18637 | 23883 | 21091 |
| *Molossus aztecus* | generalist | insectivore | 0.75 | 23 | 3691 | 3144 |
| *Molossus bondae* | generalist | insectivore | 0.77 | 1236 | 1637 | 658 |
| *Molossus coibensis* | generalist | insectivore | 0.77 | 15654 | 14718 | 11977 |
| *Molossus molossus* | generalist | insectivore | 0.67 | 26305 | 25921 | 23002 |
| *Molossus pretiosus* | generalist | insectivore | 0.77 | 3516 | 15536 | 12911 |
| *Molossus rufus* | generalist | insectivore | 0.69 | 41271 | 43375 | 42714 |
| *Molossus sinaloae* | generalist | insectivore | 0.7 | 3264 | 10000 | 9302 |
| *Mormoops megalophylla* | generalist | insectivore | 0.75 | 4568 | 7743 | 5077 |
| *Mormopterus kalinowskii* | generalist | insectivore | 0.91 | 1109 | 3026 | 2909 |
| *Myotis albescens* | generalist | insectivore | 0.63 | 24562 | 24316 | 21444 |
| *Myotis chiloensis* | generalist | insectivore | 0.88 | 1164 | 1594 | 1119 |
| *Myotis dinellii* | generalist | insectivore | 0.84 | 1659 | 4394 | 3613 |
| *Myotis izecksohni* | specialist | insectivore | 0.79 | 65 | 175 | 4 |
| *Myotis keaysi* | specialist | insectivore | 0.68 | 2992 | 11055 | 5431 |
| *Myotis levis* | specialist | insectivore | 0.88 | 620 | 287 | 7 |
| *Myotis nigricans* | generalist | insectivore | 0.66 | 40901 | 43734 | 43307 |
| *Myotis oxyotus* | generalist | insectivore | 0.77 | 3380 | 6638 | 5655 |
| *Myotis riparius* | specialist | insectivore | 0.62 | 20381 | 18502 | 8782 |
| *Myotis ruber* | specialist | insectivore | 0.83 | 753 | 339 | 7 |
| *Myotis simus* | specialist | insectivore | 0.84 | 4720 | 8589 | 6436 |
| *Natalus espiritosantensis* | specialist | insectivore | 0.8 | 80 | 29 | 44 |
| *Natalus mexicanus* | generalist | insectivore | 0.85 | 3674 | 3993 | 3686 |
| *Natalus tumidirostris* | generalist | insectivore | 0.71 | 2131 | 5972 | 5361 |
| *Noctilio albiventris* | generalist | insectivore | 0.65 | 35674 | 38618 | 37727 |
| *Noctilio leporinus* | generalist | piscivore | 0.68 | 42160 | 45146 | 44654 |
| *Nyctinomops aurispinosus* | generalist | insectivore | 0.74 | 2968 | 5418 | 8004 |
| *Nyctinomops laticaudatus* | generalist | insectivore | 0.65 | 21235 | 24485 | 21659 |
| *Nyctinomops macrotis* | generalist | insectivore | 0.72 | 23573 | 25336 | 22556 |
| *Peropteryx kappleri* | generalist | insectivore | 0.79 | 14247 | 14712 | 8715 |
| *Peropteryx leucoptera* | generalist | insectivore | 0.76 | 11029 | 8957 | 4829 |
| *Peropteryx macrotis* | generalist | insectivore | 0.67 | 36245 | 38456 | 37944 |
| *Peropteryx pallidoptera* | specialist | insectivore | 0.81 | 980 | 3512 | 2712 |
| *Phylloderma stenops* | generalist | omnivore | 0.71 | 26476 | 26756 | 18756 |
| *Phyllostomus discolor* | generalist | omnivore | 0.66 | 33896 | 33834 | 31924 |
| *Phyllostomus elongatus* | generalist | omnivore | 0.65 | 16928 | 16921 | 8877 |
| *Phyllostomus hastatus* | generalist | omnivore | 0.64 | 35102 | 35781 | 34083 |
| *Phyllostomus latifolius* | specialist | omnivore | 0.81 | 2 | 0 | 0 |
| *Platalina genovensium* | generalist | nectarivore | 0.9 | 830 | 2484 | 2437 |
| *Platyrrhinus albericoi* | specialist | frugivore | 0.81 | 1290 | 5206 | 3596 |
| *Platyrrhinus angustirostris* | specialist | frugivore | 0.82 | 895 | 4933 | 2460 |
| *Platyrrhinus aurarius* | specialist | frugivore | 0.85 | 744 | 1087 | 801 |
| *Platyrrhinus brachycephalus* | generalist | frugivore | 0.73 | 13668 | 13735 | 6908 |
| *Platyrrhinus chocoensis* | specialist | frugivore | 0.96 | 162 | 274 | 42 |
| *Platyrrhinus dorsalis* | specialist | frugivore | 0.8 | 825 | 4040 | 1952 |
| *Platyrrhinus fusciventris* | specialist | frugivore | 0.81 | 5064 | 3073 | 1853 |
| *Platyrrhinus helleri* | generalist | frugivore | 0.71 | 26763 | 29367 | 28527 |
| *Platyrrhinus incarum* | specialist | frugivore | 0.79 | 5685 | 7172 | 3445 |
| *Platyrrhinus infuscus* | generalist | frugivore | 0.76 | 2944 | 10923 | 7689 |
| *Platyrrhinus ismaeli* | generalist | frugivore | 0.86 | 256 | 911 | 132 |
| *Platyrrhinus lineatus* | specialist | frugivore | 0.7 | 2178 | 2132 | 90 |
| *Platyrrhinus masu* | specialist | frugivore | 0.85 | 474 | 3666 | 2292 |
| *Platyrrhinus nigellus* | specialist | frugivore | 0.88 | 708 | 1118 | 185 |
| *Platyrrhinus recifinus* | generalist | frugivore | 0.82 | 2190 | 3800 | 3212 |
| *Platyrrhinus vittatus* | specialist | frugivore | 0.73 | 713 | 6584 | 3564 |
| *Promops centralis* | specialist | insectivore | 0.71 | 6670 | 17449 | 9107 |
| *Pteronotus davyi* | generalist | insectivore | 0.7 | 4844 | 12809 | 5810 |
| *Pteronotus gymnonotus* | generalist | insectivore | 0.68 | 7191 | 15071 | 9386 |
| *Pteronotus personatus* | generalist | insectivore | 0.71 | 9317 | 15217 | 11316 |
| *Pteronotus rubiginosus* | specialist | insectivore | 0.71 | 10510 | 12980 | 6543 |
| *Pygoderma bilabiatum* | generalist | frugivore | 0.76 | 2468 | 3240 | 2573 |
| *Rhinophylla alethina* | specialist | frugivore | 0.89 | 191 | 211 | 38 |
| *Rhinophylla fischerae* | specialist | frugivore | 0.8 | 10789 | 10490 | 5260 |
| *Rhinophylla pumilio* | specialist | frugivore | 0.69 | 15794 | 15511 | 7650 |
| *Rhogeessa io* | generalist | insectivore | 0.68 | 8274 | 17120 | 14419 |
| *Rhogeessa minutilla* | generalist | insectivore | 0.78 | 988 | 1458 | 719 |
| *Rhynchonycteris naso* | generalist | insectivore | 0.68 | 18084 | 16434 | 7095 |
| *Saccopteryx bilineata* | generalist | insectivore | 0.69 | 18698 | 17991 | 9271 |
| *Saccopteryx canescens* | generalist | insectivore | 0.77 | 14905 | 14231 | 11596 |
| *Saccopteryx leptura* | generalist | insectivore | 0.66 | 21012 | 20895 | 18173 |
| *Scleronycteris ega* | generalist | insectivore | 0.86 | 2432 | 6063 | 5293 |
| *Sphaeronycteris toxophyllum* | specialist | frugivore | 0.73 | 7610 | 7645 | 4052 |
| *Sturnira aratathomasi* | specialist | frugivore | 0.93 | 126 | 720 | 120 |
| *Sturnira bidens* | specialist | frugivore | 0.82 | 676 | 734 | 55 |
| *Sturnira bogotensis* | specialist | frugivore | 0.79 | 484 | 849 | 160 |
| *Sturnira erythromos* | specialist | frugivore | 0.72 | 1209 | 3120 | 780 |
| *Sturnira lilium* | specialist | frugivore | 0.76 | 1603 | 1684 | 87 |
| *Sturnira ludovici* | specialist | frugivore | 0.74 | 1115 | 11273 | 6860 |
| *Sturnira luisi* | specialist | frugivore | 0.84 | 418 | 713 | 200 |
| *Sturnira magna* | specialist | frugivore | 0.74 | 2873 | 3778 | 2118 |
| *Sturnira oporaphilum* | specialist | frugivore | 0.76 | 897 | 1133 | 242 |
| *Sturnira tildae* | specialist | frugivore | 0.65 | 16481 | 15533 | 7555 |
| *Tadarida brasiliensis* | generalist | insectivore | 0.71 | 23595 | 33340 | 32446 |
| *Thyroptera discifera* | generalist | insectivore | 0.74 | 11999 | 6314 | 4306 |
| *Thyroptera tricolor* | generalist | insectivore | 0.7 | 23171 | 25461 | 18028 |
| *Tonatia bidens* | specialist | omnivore | 0.82 | 1090 | 1616 | 86 |
| *Tonatia saurophila* | generalist | insectivore | 0.76 | 14035 | 18601 | 16740 |
| *Trachops cirrhosus* | generalist | carnivore | 0.67 | 22182 | 20458 | 17622 |
| *Trinycteris nicefori* | generalist | frugivore | 0.75 | 9225 | 11788 | 9902 |
| *Uroderma bilobatum* | generalist | frugivore | 0.64 | 27404 | 28749 | 27700 |
| *Uroderma magnirostrum* | specialist | frugivore | 0.69 | 17345 | 15711 | 6937 |
| *Vampyressa melissa* | specialist | frugivore | 0.83 | 993 | 4640 | 2933 |
| *Vampyressa thyone* | specialist | frugivore | 0.71 | 8170 | 16273 | 9003 |
| *Vampyriscus bidens* | specialist | frugivore | 0.75 | 15179 | 13126 | 6465 |
| *Vampyriscus brocki* | specialist | frugivore | 0.78 | 2146 | 3488 | 2158 |
| *Vampyriscus nymphaea* | specialist | frugivore | 0.85 | 404 | 482 | 118 |
| *Vampyrodes caraccioli* | generalist | frugivore | 0.72 | 16653 | 16130 | 13190 |
| *Vampyrodes major* | generalist | frugivore | 0.75 | 1504 | 2482 | 2257 |
| *Vampyrum spectrum* | specialist | carnivore | 0.68 | 8625 | 15145 | 7576 |

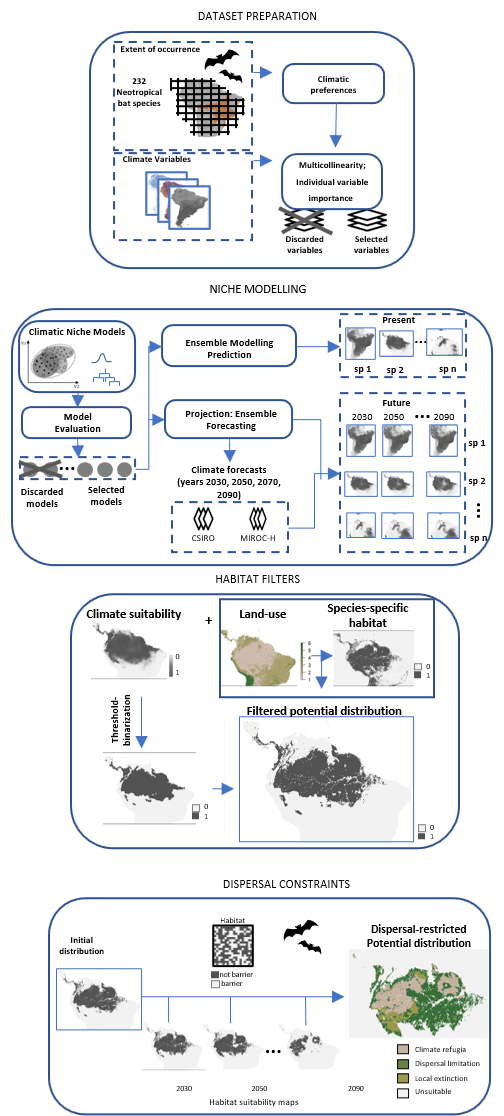

**Appendix S3:** Methods workflow used to forecast the combined effects of climate and land use change on Neotropical bats biodiversity. Figure adapted from Sales et al. (2020).

**Amazon**

**Appendix S4:** Restructuring of Neotropical bats diversity in the Amazon under the business-as-usual (B.A.U) scenario of climate and land-use changes and assuming limited dispersal (bat species have the ability to disperse only over analog environments) Frug = frugivore, Nect = nectarivore, Inse = insectivores, Pisc = piscivores, Sang = sanguivores, Carn = carnivores and Omni = omnivore. Contraction = range contraction, Expansion = increase range size.

|  | Phytophagous | |  | Animalivorous | | | |  | Omnivores |  | Total |
| --- | --- | --- | --- | --- | --- | --- | --- | --- | --- | --- | --- |
| Amazon |  |  |  |  |  |  |  |  |  |  |  |
|  | Frug | Nect |  | Inse | Pisc | Sang | Carn |  | Omni |  |  |
| Contraction | 44 | 8 |  | 54 |  | 2 | 3 |  | 4 |  | 115 |
| Expansion | 26 | 18 |  | 50 | 1 | 1 |  |  | 1 |  | 97 |
| Extinct |  |  |  |  |  |  |  |  |  |  |  |
| Gains |  |  |  |  |  |  |  |  |  |  |  |
| Total | 70 | 26 |  | 104 | 1 | 3 | 3 |  | 5 |  |  |

**Atlantic Forest**

**Appendix S5:** Restructuring of Neotropical bat diversity in the Atlantic Forest under the business-as-usual (B.A.U) scenario of climate and land-use changes and assuming limited dispersal (bat species have the ability to disperse only over analog environments) Frug = frugivore, Nect = nectarivore, Inse = insectivores, Pisc = piscivores, Sang = sanguivores, Carn = carnivores and Omni = omnivore. Contraction = range contraction, Expansion = increase range size, Extinct (no analog environments in the future) and Gain (bat species on the move from diferent biogeographyc region).

| Atlantic | Phytophagous | |  | Animalivorous | | | |  | Omnivores |  | Total |
| --- | --- | --- | --- | --- | --- | --- | --- | --- | --- | --- | --- |
| forest | Frug | Nect |  | Inse | Pisc | Sang | Carn |  | Omni |  |  |
| Contraction | 10 | 1 |  | 26 |  | 2 | 1 |  | 2 |  | 42 |
| Expansion | 4 | 3 |  | 26 | 1 | 1 | 1 |  | 3 |  | 39 |
| Extinct | 5 | 3 |  | 7 |  |  |  |  |  |  | 15 |
| Gains |  | 1 |  | 2 |  |  |  |  |  |  | 3 |
| Total | 19 | 8 |  | 61 | 1 | 3 | 2 |  | 5 |  |  |

**Caatinga**

**Appendix S6:** Restructuring of Neotropical bat diversity in the Caatinga under the business-as-usual (B.A.U) scenario of climate and land-use changes and assuming limited dispersal (bat species have the ability to disperse only over analog environments) Frug = frugivore, Nect = nectarivore, Inse = insectivores, Pisc = piscivores, Sang = sanguivores, Carn = carnivores and Omni = omnivore. Contraction = range contraction, Expansion = increase range size, Extinct (no analog environments in the future) and Gain (bat species on the move from diferent ecoregion).

|  | Phytophagous | |  | Animalivorous | | | |  | Omnivores | Total |
| --- | --- | --- | --- | --- | --- | --- | --- | --- | --- | --- |
| Caatinga |  |  |  |  |  |  |  |  |  |  |
|  | Frug | Nect |  | Inse | Pisc | Sang | Carn |  | Omni |  |
| Contraction | 8 | 2 |  | 37 | 1 | 2 | 2 |  | 4 | 56 |
| Expansion | 3 | 1 |  | 1 |  | 1 |  |  |  | 6 |
| Extinct | 6 | 1 |  | 9 |  |  |  |  | 1 | 17 |
| Gains | 1 | 1 |  | 4 |  |  |  |  |  | 6 |
| Total | 18 | 5 |  | 51 | 1 | 3 | 2 |  | 5 |  |

**West indies**

**Appendix S7:** Restructuring of Neotropical bat diversity in the West indies under the business-as-usual (B.A.U) scenario of climate and land-use changes and assuming limited dispersal (bat species have the ability to disperse only over analog environments) Frug = frugivore, Nect = nectarivore, Inse = insectivores, Pisc = piscivores, Sang = sanguivores, Carn = carnivores and Omni = omnivore. Contraction = range contraction, Expansion = increase range size, Extinct (no analog environments in the future) and Gain (bat species on the move from diferent ecoregion).

|  | Phytophagous | |  | Animalivorous | | | |  | Omnivores | Total |
| --- | --- | --- | --- | --- | --- | --- | --- | --- | --- | --- |
| West indies |  |  |  |  |  |  |  |  |  |  |
|  | Frug | Nect |  | Inse | Pisc | Sang | Carn |  | Omni |  |
| Contraction | 2 | 1 |  | 10 | 1 |  |  |  |  | 14 |
| Expansion |  |  |  |  |  |  |  |  |  |  |
| Extinct |  | 3 |  | 3 |  |  |  |  | 1 | 7 |
| Gains |  |  |  |  |  |  |  |  |  |  |
| Total | 2 | 4 |  | 13 | 1 |  |  |  | 1 |  |

**Cerrado and Chaco**

**Appendix S8:** Restructuring of Neotropical bat diversity in the Cerrado and Chaco under the business-as-usual (B.A.U) scenario of climate and land-use changes and assuming limited dispersal (bat species have the ability to disperse only over analog environments) Frug = frugivore, Nect = nectarivore, Inse = insectivores, Pisc = piscivores, Sang = sanguivores, Carn = carnivores and Omni = omnivore. Contraction = range contraction, Expansion = increase range size, Extinct (no analog environments in the future) and Gain (bat species on the move from diferent ecoregion).

| Cerrado | Phytophagous | |  | Animalivorous | | | |  | Omnivores | Total |
| --- | --- | --- | --- | --- | --- | --- | --- | --- | --- | --- |
| and |  |  |  |  |  |  |  |  |  |  |
| Chaco | Frug | Nect |  | Inse | Pisc | Sang | Carn |  | Omni |  |
| Contraction | 28 | 5 |  | 35 |  | 2 | 3 |  | 2 | 75 |
| Expansion | 8 | 5 |  | 26 | 1 | 1 |  |  | 3 | 44 |
| Extinct | 5 | 1 |  | 15 |  |  |  |  |  | 21 |
| Gains | 2 | 4 |  | 7 |  |  |  |  | 1 | 14 |
| Total | 43 | 15 |  | 83 | 1 | 3 | 3 |  | 6 |  |

**Dry Northern South America**

**Appendix S9:** Restructuring of Neotropical bat diversity in the Dry Northern South America under the business-as-usual (B.A.U) scenario of climate and land-use changes and assuming limited dispersal (bat species have the ability to disperse only over analog environments) Frug = frugivore, Nect = nectarivore, Inse = insectivores, Pisc = piscivores, Sang = sanguivores, Carn = carnivores and Omni = omnivore. Contraction = range contraction, Expansion = increase range size, Extinct (no analog environments in the future) and Gain (bat species on the move from diferent ecoregion).

| Dry | Phytophagous | |  | Animalivorous | | | |  | Omnivores | Total |
| --- | --- | --- | --- | --- | --- | --- | --- | --- | --- | --- |
| Northern | Frug | Nect |  | Inse | Pisc | Sang | Carn |  | Omni |  |
| Contraction | 29 | 11 |  | 67 | 1 | 3 | 3 |  | 4 | 118 |
| Expansion | 9 | 6 |  | 22 |  |  |  |  |  | 37 |
| Extinct | 6 | 2 |  | 6 |  |  |  |  | 1 | 15 |
| Gains | 1 | 1 |  | 1 |  |  |  |  |  | 3 |
| Total | 45 | 20 |  | 96 | 1 | 3 | 3 |  | 5 |  |

**Dry Western South America**

**Appendix S10:** Restructuring of Neotropical bat diversity in the Dry Western South America under the business-as-usual (B.A.U) scenario of climate and land-use changes and assuming limited dispersal (bat species have the ability to disperse only over analog environments) Frug = frugivore, Nect = nectarivore, Inse = insectivores, Pisc = piscivores, Sang = sanguivores, Carn = carnivores and Omni = omnivore. Contraction = range contraction, Expansion = increase range size, Extinct (no analog environments in the future) and Gain (bat species on the move from diferent ecoregion).

| Dry | Phytophagous | |  | Animalivorous | | | |  | Omnivores |  | Total |
| --- | --- | --- | --- | --- | --- | --- | --- | --- | --- | --- | --- |
| Western | Frug | Nect |  | Inse | Pisc | Sang | Carn |  | Omni |  |  |
| Contraction | 15 | 2 |  | 18 |  | 1 | 1 |  |  |  | 37 |
| Expansion | 12 | 12 |  | 26 | 1 | 1 | 2 |  | 3 |  | 57 |
| Extinct | 6 |  |  | 2 |  |  |  |  |  |  | 8 |
| Gains | 19 | 5 |  | 34 |  | 1* |  |  | 2 |  | 61 |
| Total | 27 | 14 |  | 44 | 1 | 2 | 3 |  | 3 |  |  |

******Diaemus youngi*

**Mesoamerica**

**Appendix S11:** Restructuring of Neotropical bat diversity in the Mesoamerica under the business-as-usual (B.A.U) scenario of climate and land-use changes and assuming limited dispersal (bat species have the ability to disperse only over analog environments) Frug = frugivore, Nect = nectarivore, Inse = insectivores, Pisc = piscivores, Sang = sanguivores, Carn = carnivores and Omni = omnivore. Contraction = range contraction, Expansion = increase range size, Extinct (no analog environments in the future) and Gain (bat species on the move from diferent ecoregion).

|  | Phytophagous | |  | Animalivorous | | | |  | Omnivores | Total |
| --- | --- | --- | --- | --- | --- | --- | --- | --- | --- | --- |
| Mesoamerica |  |  |  |  |  |  |  |  |  |  |
|  | Frug | Nect |  | Inse | Pisc | Sang | Carn |  | Omni |  |
| Contraction | 18 | 5 |  | 28 |  | 2 | 3 |  | 3 | 59 |
| Expansion | 4 | 4 |  | 35 | 1 | 1 |  |  | 1 | 46 |
| Extinct |  |  |  |  |  |  |  |  |  |  |
| Gains | 13 | 4 |  | 11 |  |  |  |  |  | 28 |
| Total | 35 | 13 |  | 74 | 1 | 3 | 3 |  | 4 |  |

**Andean grasslands**

**Appendix S12:** Restructuring of Neotropical bat diversity in the Andean grasslands under the business-as-usual (B.A.U) scenario of climate and land-use changes and assuming limited dispersal (bat species have the ability to disperse only over analog environments) Frug = frugivore, Nect = nectarivore, Inse = insectivores, Pisc = piscivores, Sang = sanguivores, Carn = carnivores and Omni = omnivore. Contraction = range contraction, Expansion = increase range size, Extinct (no analog environments in the future) and Gain (bat species on the move from diferent ecoregion).

| Andean | Phytophagous | |  | Animalivorous | | | |  | Omnivores | Total |
| --- | --- | --- | --- | --- | --- | --- | --- | --- | --- | --- |
| grasslands | Frug | Nect |  | Inse | Pisc | Sang | Carn |  | Omni |  |
| Contraction | 25 | 9 |  | 41 |  | 1 | 1 |  | 1 | 78 |
| Expansion | 10 | 11 |  | 30 | 1 | 2 | 2 |  | 3 | 59 |
| Extinct | 16 | 2 |  | 3 |  |  |  |  |  | 21 |
| Gains | 2 | 1 |  | 5 |  |  |  |  |  | 8 |
| Total | 53 | 23 |  | 79 | 1 | 3 | 3 |  | 4 |  |

**Patagonian steppe**

**Appendix S13:** Restructuring of Neotropical bat diversity in the Patagonian steppe under the business-as-usual (B.A.U) scenario of climate and land-use changes and assuming limited dispersal (bat species have the ability to disperse only over analog environments) Frug = frugivore, Nect = nectarivore, Inse = insectivores, Pisc = piscivores, Sang = sanguivores, Carn = carnivores and Omni = omnivore. Contraction = range contraction, Expansion = increase range size, Extinct (no analog environments in the future) and Gain (bat species on the move from diferent ecoregion).

| Patagonian | Phytophagous | |  | Animalivorous | | | |  | Omnivores | Total |
| --- | --- | --- | --- | --- | --- | --- | --- | --- | --- | --- |
| steppe | Frug | Nect |  | Inse | Pisc | Sang | Carn |  | Omni |  |
| Contraction |  |  |  | 2 |  | 1 |  |  |  | 3 |
| Expansion | 1 |  |  | 18 | 1 |  |  |  |  | 20 |
| Extinct | 4 |  |  | 3 |  |  |  |  |  | 7 |
| Gains | 3 |  |  | 5 |  | 1* | 1 |  |  | 10 |
| Total | 8 |  |  | 28 | 1 | 2 | 1 |  |  |  |

**Diphylla ecaudata*

IPCC, 2014. Climate Change 2014: Synthesis Report. Contribution of Working Groups I, II and III to the Fifth Assessment Report of the Intergovernmental Panel on Climate Change (RK Pachauri and L Meyer, Eds.).

IPCC, 2019: Climate Change and Land: an IPCC special report on climate change,desertification, land degradation, sustainable land management, food security,and greenhouse gas fluxes in terrestrial ecosystems. [Shukla, P.R., Skea, J.,Calvo Buendia, E., Masson-Delmotte, V., Pörtner, H.-O., Roberts, D.C., Zhai, P.,Slade, R., Connors, S., van Diemen, R., Ferrat, M., Haughey, E., Luz, S., Neogi, S.,Pathak, M., Petzold, J., Portugal Pereira, J., Vyas, P., Huntley, E., Kissick, K.,Belkacemi, M., Malley, J. (eds.)]. Retrieved from www.ipcc.ch.

Kriticos, D.J., Webber, B.L., Leriche, A., Ota, N., Macadam, I., Bathols, J., Scott, J.K., 2012. CliMond: global high-resolution historical and future scenario climate surfaces for bioclimatic modelling. Methods Ecol. Evol. 3, 53–64. https://doi.org/10.1111/j.2041-210X.2011.00134.x

Li, X., Chen, G., Liu, X., Liang, X., Wang, S., Chen, Y., Pei, F., Xu, X., 2017. A New Global Land-Use and Land-Cover Change Product at a 1-km Resolution for 2010 to 2100 Based on Human–Environment Interactions. Ann. Am. Assoc. Geogr. 107, 1040–1059. https://doi.org/10.1080/24694452.2017.1303357

Marquaridt, D.W., 1970. Generalized Inverses, Ridge Regression, Biased Linear Estimation, and Nonlinear Estimation. Technometrics 12, 591–612. https://doi.org/10.1080/00401706.1970.1048869

Mickleburgh, S.P., Hutson, A.M., Racey, P.A., 2002. A review of the global conservation status of bats. Oryx 36, 18–34. https://doi.org/DOI: 10.1017/S0030605302000054

Morrone, J.J., 2014. Biogeographical regionalisation of the Neotropical region. Zootaxa 3782(1), 1-110.

Naimi, B., Hamm, N.A.S., Groen, T.A., Skidmore, A.K., Toxopeus, A.G., 2014. Where is positional uncertainty a problem for species distribution modelling? Ecography (Cop.). 37, 191–203. https://doi.org/10.1111/j.1600-0587.2013.00205.x

New, M., Lister, D., Hulme, M., Makin, I., 2002. A high-resolution data set of surface climate over global land areas. Climate Research 21:1–25.

Olson, D.M., Dinerstein, E., Wikramanayake, E.D., Burgess, N.D., Powell, G.V.N., Underwood, E.C., D’amico, J.A., Itoua, I., Strand, H.E., Morrison, J.C., Loucks, C.J., Allnutt, T.F., Ricketts, T.H., Kura, Y., Lamoreux, J.F., Wettengel, W.W., Hedao, P., Kassem, K.R., 2001. Terrestrial Ecoregions of the World: A New Map of Life on Earth: A new global map of terrestrial ecoregions provides an innovative tool for conserving biodiversity. Bioscience 51, 933–938. https://doi.org/10.1641/0006-3568(2001)051[0933:TEOTWA]2.0.CO;2

Ribeiro, B.R., Sales, L.P., De Marco Jr., P., Loyola, R., 2016. Assessing Mammal Exposure to Climate Change in the Brazilian Amazon. PLoS One 11, 1–13. https://doi.org/10.1371/journal.pone.0165073

Sales, L.P., Ribeiro, B.R., Pires, M.M., Chapman, C.A., Loyola, R., 2019. Recalculating route: dispersal constraints will drive the redistribution of Amazon primates in the Anthropocene. Ecography (Cop.). 42, 1789–1801. https://doi.org/10.1111/ecog.04499

Sales, L.P., Galetti, M., Pires, M.M., 2020. Climate and land-use change will lead to a faunal “savannization” on tropical rainforests. Glob. Chang. Biol. 26, 7036–7044. https://doi.org/10.1111/gcb.15374

Sherwin, H.A., Montgomery, W.I., Lundy, M.G., 2013. The impact and implications of climate change for bats. Mamm. Rev. 43, 171–182. https://doi.org/10.1111/j.1365-2907.2012.00214.x

Simmons, N.B., 2005. Order_Chiroptera_T.E.Lawlor.pdf, in: REEDER, D.E.W.and D.M. (Ed.), Mammals Species of the World: A Taxonomic and Geographic Reference. Johns Hopkins University Press, Baltimore, pp. 312–529.

Vilhena, D.A., Antonelli, A., 2015. A network approach for identifying and delimiting biogeographical regions. Nat. Commun. 6, 6848. https://doi.org/10.1038/ncomms7848

Combined impacts of climate and land use change and the future restructuring of Neotropical bat biodiversity

Fernando Gonçalves, Lilian P. Sales, Mauro Galetti, Mathias M. Pires

31/07/2021

Code by Lilian P. Sale

CLIMATE

Climond Bioclim data present and furute (only year 2030 is shown)

```
devtools::install_github('babaknaimi/sdm') # to install the latest version of the sdm package from github
library(sdm)
library(spThin )
library(dismo)
library(rgbif)
library(scrubr)
library(spocc)
library(rvertnet)
library(plyr)
library(scrubr)
library(usdm)
library(rgdal)
library(sp)
library(spatialEco)
library(plyr)
library(letsR)

setwd("D:/Leddiv/Climate/Climond_10'/CM10_1975H_Bio_ASCII_V1.2/CM10_1975H_Bio_V1.2")
pres <- list.files(pattern = ".txt")
pres <- pres[1:19]
pres <- stack(pres)
pres@crs <- CRS("+proj=longlat +datum=WGS84")

# 2030 -----------------------------------------------------------------------------------------

setwd("D:/Leddiv/Climate/Climond_10'/CM10_2030_A1B_CS_Bio_ASCII_V1.2/CM10_2030_A1B_CS_Bio_V1.2")
f30a <- list.files(pattern = ".txt")
f30a <- stack(f30a[1:19])
f30a@crs <- CRS("+proj=longlat +datum=WGS84")

setwd("D:/Leddiv/Climate/Climond_10'/CM10_2030_A1B_MR_Bio_ASCII_V1.2/CM10_2030_A1B_MR_Bio_V1.2")
f30b <- list.files(pattern = ".txt")
f30b <- stack(f30b[1:19])
f30b@crs <- CRS("+proj=longlat +datum=WGS84")

setwd("D:/Leddiv/Climate/Climond_10'/CM10_2030_A2_CS_Bio_ASCII_V1.2/CM10_2030_A2_CS_Bio_V1.2")
f30c <- list.files(pattern = ".txt")
f30c <- stack(f30c[1:19])
f30c@crs <- CRS("+proj=longlat +datum=WGS84")

setwd("D:/Leddiv/Climate/Climond_10'/CM10_2030_A2_MR_Bio_ASCII_V1.2/CM10_2030_A2_MR_Bio_V1.2")
f30d <- list.files(pattern = ".txt")
f30d <- stack(f30d[1:19])
f30d@crs <- CRS("+proj=longlat +datum=WGS84")

# Check for name consistency
names(f30a) <- names(pres)
names(f30b) <- names(pres)
names(f30c) <- names(pres)
names(f30d) <- names(pres)
```

#### SPECIES DATA

Select South American bats and environmental info from IUCN range maps

```
memory.limit() #Allow greater use of memory
Mam <- spTransform(Mam, CRS("+proj=longlat +datum=WGS84"))

#Pres/Abs matrix
presab_poligono <- lets.presab (Mam, xmn=-131,xmx=-35,ymn=-56, ymx=66,
                                resol=0.16, count=T, crs = CRS("+proj=longlat +datum=WGS84"),
                                cover=0.5, remove.cells=T)

# Plotting the raster (a species richness map)
plot(presab_poligono[[2]])
setwd("D:/Leddiv/lilian/Bats")
save.image("PresAb.RData")

#load("PresAb.RData")

# Getting the pres/abs matrix
matrizPA <- as.data.frame(presab_poligono[[1]])
str(matrizPA)

#str(matrizPA)
#write.csv(matrizPA, "matrizPA.csv")
# Create Presences file
#install.packages("tidyverse", dep=T)
library(tidyverse)
library(reshape2)

a <- data.frame(matrizPA)
b <- melt(a, id.vars = c("Longitude.x.", "Latitude.y."))
head(b)
ocorr <- b %>%
  filter(value > 0)
ocorr <- as.data.frame(ocorr)
ocorr <- ocorr[,c("Longitude.x.","Latitude.y.","variable")]
colnames(ocorr) <- c("lon","lat","variable")
head(ocorr)
tail(ocorr)

.rs.unloadPackage("tidyr") # Como este pacote contém funções que são usadas em outros pacotes, estava dando erro

# Remove species with less than 30 records
occ_count <- table(unlist(ocorr$variable))
ocorr <- ocorr[ocorr$variable %in% names(occ_count)[occ_count>30],]
occ_count1 <- occ_count[occ_count>30]

write.csv(occ_count1, "D:/Leddiv/lilian/Bats/contagem_ocorr.csv")

sp_names <- unique(ocorr$variable)

resu <- matrix(nrow = length(sp_names), ncol = 15)
colnames(resu) <- c("sp_name","records","AUC","COR","Deviance","TSS", "iniDist", "30A1B", "30A2", "50A1B", "50A2", "70A1B", "70A2", "90A1B", "90A2")
vars <- NULL
```

#### Ecologinal niche modelling

Using ensembles and IUCN range maps

```
for (i in 1:length (sp_names)){

  t <- sp_names[[i]]
  sp <- ocorr[ocorr$variable==t,]

if(nrow(sp)<100){
  x = 1
}

if(nrow(sp)>100){
  x= 0.5}

if(nrow(sp)>500){
  x= 0.25}

if(nrow(sp)>1000){
  x= 0.125}

  spSample <- sample(1:nrow(sp), replace=F, size=round(x * nrow(sp)))

  sp <-sp[spSample ,c("lon", "lat")]
  sp$species <- 1

  coordinates(sp) <- ~ lon + lat
  sp@proj4string <- CRS("+proj=longlat +datum=WGS84")
  plot(sp, main = t)

  bb <- bbox(sp)

  bb.buf <- extent(bb[1]-10, bb[3]+10, bb[2]-10, bb[4]+10)

  envs.backg <- crop(pres, bb.buf)

  # Remove collinear variables

  spx <- extract(envs.backg, sp) # extract from file
  spx <- data.frame(spx) #convert to dataframe

  v <- vifcor(spx, th=0.7)      # check collinearity (variance inflation and correlation)
  bio_i <- exclude(envs.backg, v) # exclude collinear predictors

  # generate sdmData
  d <- sdm::sdmData(species~ ., train=sp, predictors= bio_i, bg=list(n=10000,method='gRandom',remove=TRUE))

  # generate sdm model
  m <- sdm(species ~ ., d, methods=c("brt","maxlike"),
           replication=c("sub"), test.p=25, n=100,
           parallelSettings = list(ncore=10, method='parallel'))

 # m

  # Ensembling Present
  en <- ensemble(m, bio_i,
                 setting=list(method='weighted',stat='TSS'),
                 parallelSettings = list(ncore=10, method='parallel'))

  plot(en, main="Present")

  writeRaster(en,  paste0('D:/Leddiv/lilian/Bats/Mapas/', gsub(" ", "_", t),"_pres.tif"), format = "GTiff", overwrite=TRUE)

  # Evaluation
  e <- getEvaluation(m)

  d1 <- as.data.frame(d)

  # Save which variables were used and evaluation results for all species
  resu[i, "sp_name"] <- gsub(" ", "_", t)
  resu[i, "records"] <- nrow(d1[d1$species>0, ])
  resu[i, "AUC"] <- round(mean(e$AUC), 2)
  resu[i, "COR"] <- round(mean(e$COR), 2)
  resu[i, "Deviance"] <- round(mean(e$Deviance),2)
  resu[i, "TSS"] <- round(mean(e$TSS),2)

  vars <- c(as.character(t), names(bio_i), vars)

  # Find binarization threshold
  df <- data.frame(as.data.frame(d),coordinates(d)) # presence points and predictors associated
  pr <- extract(en, df[,c("lon","lat")])

  ev <- evaluates(df$species, pr) # evaluate prediction (observed vs expected)
  th <- ev@threshold_based$threshold[[2]] # threshold that maximizes sensitiv + specificity

  # Binary prediction
  pa <- en
  pa[] <- ifelse(pa[] >= th, 1, 0) # convert from continuous to binary

  plot(pa, main = t)

  # present PA
  writeRaster(pa,   format = "GTiff",
              paste0("D:/Leddiv/lilian/Bats/Mapas/",gsub(" ", "_", t),"_pres_PA.tif"),
              overwrite = T)

  iniDist <- length(pa[pa==1])

  # Ensembling Future
  # 2030 - A1B
    # rcp8.5
  enf1 <- ensemble(m, crop(f30a, bb.buf),
                   setting=list(method='weighted',stat='TSS'),
                   parallelSettings = list(ncore=10, method='parallel'))

  enf2 <- ensemble(m,  crop(f30b, bb.buf),
                   setting=list(method='weighted',stat='TSS'),
                   parallelSettings = list(ncore=10, method='parallel'))

  # 2030 - A2
  enf3 <- ensemble(m,  crop(f30c, bb.buf),
                   setting=list(method='weighted',stat='TSS'),
                   parallelSettings = list(ncore=10, method='parallel'))

  enf4 <- ensemble(m,  crop(f30d, bb.buf),
                   setting=list(method='weighted',stat='TSS'),
                   parallelSettings = list(ncore=10, method='parallel'))

  enf30.1 <- mean(enf1, enf2)
  #plot(enf30.1, main = t)

  enf30.2 <- mean(enf3, enf4)
  #plot(enf30.2, main = t)

  writeRaster(enf30.1,  paste0("D:/Leddiv/lilian/Bats/Mapas/", gsub(" ", "_", t),"_2030_A1B.tif"), format = "GTiff", overwrite=TRUE)
  writeRaster(enf30.2,  paste0("D:/Leddiv/lilian/Bats/Mapas/", gsub(" ", "_", t),"_2030_A2.tif"), format = "GTiff", overwrite=TRUE)

  # Binary prediction future
  #A1B
  paf1 <- enf30.1
  paf1[] <- ifelse(paf1[] >= th, 1, 0) # convert from continuous to binary

  #plot(paf1, main=paste(t, "A1B"))

  finalDist1 <- length(paf1[paf1==1])

  # Future PA
  writeRaster(paf1,   format = "GTiff",
              paste0("D:/Leddiv/lilian/Bats/Mapas/",gsub(" ", "_", t),"_2030_PA_A1B.tif"),
              overwrite = T)

  resu[i, "30A1B"] <- finalDist1

  # A2igation
  paf2 <- enf30.2
  paf2[] <- ifelse(paf2[] >= th, 1, 0) # convert from continuous to binary

  plot(paf2, main=paste(t, "A2"))

  finalDist2 <- length(paf2[paf2==1])

  # Future PA
  writeRaster(paf2,   format = "GTiff",
              paste0("D:/Leddiv/lilian/Bats/Mapas/",gsub(" ", "_", t),"_2030_PA_A2.tif"),
              overwrite = T)

  resu[i, "30A2"] <- finalDist2

}

write.csv(resu, "D:/Leddiv/lilian/Bats/Resu/resu.csv")
write.csv(vars, 'D:/Leddiv/lilian/Bats/Resu/vars.csv')
```

Now, we will apply different thresholds of forest cover, depending on species vulnerability to forest loss. Three functional groups are defined: - extreme forest specialists - moderate forest specialists - generalists

For the extreme forest specialists, suitable climate regions with less than 50% forest cover are considered unable to support viable populations.

```
library(rgeos)

# ------ Extreme forest specialists (threatened habitat specialists)
# Species assessed as Critically Endangered (CR), Endangered (EN), or Vulnerable (VU) are referred to as "threatened" species
# remove predicted presences from landscapes with less than 50% tree cover

iu <- read.csv("C:/Users/lilia/OneDrive/Pos-doc/PNPD/Bats/UFO/Planilhas/Lista_sp.csv", h=T) # List with information on IUCN status
th <- c("CR", "EN", "VU")
th_sp <- iu[iu$IUCN %in% th, ] # select only threatened by IUCN
sp <- as.character(th_sp$Species) # these species will be removed from landscapes with less than 50% tree cover
tableS1 <- data.frame(cbind(sp, rep("Extreme forest specialist", length(sp)))) # To incorporate this information on the Table S1
colnames(tableS1) <- c("Species", "Classification")
sp <- gsub(" ", "_", sp)

# Landscapes with less than 50% tree cover - do not provide habitat for extreme habitat specialists
land50 <- ref
land50[land50 < 50] <- 0
land50[land50 >=50 ] <- 1

land50 <- rasterToPolygons(land50, n=16, na.rm=TRUE, dissolve=T) # create mask with only landscapes of more than 50% forest cover

library(maptools)
writeSpatialShape(land50, "C:/Users/lilia/OneDrive/Pos-doc/PNPD/Bats/UFO/Defor/land50.shp")

land50 <- readOGR("C:/Users/lilia/OneDrive/Pos-doc/PNPD/Bats/UFO/Defor/land50.shp")

# Now remove threatened species from outside these areas

setwd("C:/Users/lilia/OneDrive/Pos-doc/PNPD/Bats/UFO/Mapas")
dir()

  for (i in 1: length(sp)) {

    list1 <- list.files(pattern=paste0(sp[i], "_PA_"))

        for (j in 1: length(list1)){

          a <- raster(list1[j])
          b <- (mask(a, land50))
          writeRaster(b, paste0("C:/Users/lilia/OneDrive/Pos-doc/PNPD/Bats/UFO/Mapas_Defaun/", list1[j]), format = "GTiff", overwrite = T)
      }
  }
```

Same same procedure is applied to moderate forest specialists, but to a 30% forest cover threshold.

```
# ------- Moderate forest specialists (not threatened)
# remove predicted presences from landscapes with less than 30% tree cover

no_th <- c("LC", "NT")
no_th_sp <- iu[iu$IUCN %in% no_th, ] # select only non-threatened

sp_sp <- no_th_sp[no_th_sp$Specialists==1, ] # select only specialists non-threatened
sp_sp <- as.character(sp_sp$Species) # these species will be removed from landscapes with less than 50% tree cover
tableS1.1 <- data.frame(cbind(sp_sp, rep("Moderate forest specialist", length(sp_sp)))) # To incorporate this information on the Table S1
colnames(tableS1.1) <- c("Species", "Classification")
sp_sp <- gsub(" ", "_", sp_sp)

# Landscapes with more than 30% tree cover - still may provide habitat for habitat specialists
land30 <- ref
land30[land30 < 30] <- 0
land30[land30 >=30 ] <- 1
plot(land30)

land30 <- rasterToPolygons(land30, n=16, na.rm=TRUE, dissolve=T) # create mask with only landscapes of more than 50% forest cover
writeSpatialShape(land30, "C:/Users/lilia/OneDrive/Pos-doc/PNPD/Bats/UFO/Defor/land30.shp")
plot(land30)
land30 <- readOGR("C:/Users/lilia/OneDrive/Pos-doc/PNPD/Bats/UFO/Defor/land30.shp")

# Now remove specialist species from outside these areas
setwd("C:/Users/lilia/OneDrive/Pos-doc/PNPD/Bats/UFO/Mapas")
dir()

for (i in 1: length(sp_sp)) {

  list1 <- list.files(pattern=paste0(sp_sp[i], "_PA_"))

  for (j in 1: length(list1)){

    a <- raster(list1[j])
    b <- (mask(a, land30))
    writeRaster(b, paste0("C:/Users/lilia/OneDrive/Pos-doc/PNPD/Bats/UFO/Mapas_Defaun/", list1[j]), format = "GTiff", overwrite = T)
  }
}
```

### Dispersal-restricted potential distribution of bats

In addition to climate suitability , we also simulate occupancy of potential suitable future areas under dispersal constraints posed by landscape fragmentation. The absence of trees was considered a strong barrier to extreme forest specialists; a weak barrier to moderate forest specialists; not a barrier to generalists

```
library(raster)
library(rgdal)
library(MigClim)
library(sp)

barrier <- tree # raster with future forest cover projections
resu <- NULL # Table to store results

# Extreme habitat specialists
# Barrier type = "strong"

for (i in 1: length(ext)) {

  setwd("D:/Leddiv/lilian/Bats/Disp_bar")

  a <- list.files(pattern = paste0(ext[i]))

  iniDist <- raster(paste0("D:/Leddiv/lilian/Bats/Mapas_Defaun/", ext[i], "_pres_PA.tif"))

  barr <- crop(barrier, extent(iniDist)) # species-specific extent of barrier cell
  barr <- as.data.frame(barr, xy=F)
  barr[is.na(barr)] <- 0

  iniDist <- as.data.frame(iniDist, xy=T)
  iniDist[is.na(iniDist)] <- 0

 # rcp45
    setwd("D:/Leddiv/lilian/Bats/Disp_bar")
    b <- stack(a)
    b45 <- raster::subset(b, grep("rcp45", names(b), value=T))
    b45@crs <- barrier@crs

    hsMap <- as.data.frame(b45)   # get coordinates

    hsMap[is.na(hsMap)] <- 0      # all NAs converted to 0

  ### Dispersal

  MigClim.migrate(iniDist = iniDist,
                  hsMap = hsMap,
                  rcThreshold = 1, #artificial threshold, binarization was made during SDM
                  envChgSteps = ncol(hsMap), dispSteps = ncol(hsMap),
                  barrier = barr, barrierType = "strong", # Barrier is strong to extreme habitat specialists
                  replicateNb=3,
                  testMode=FALSE,
                  fullOutput=T, keepTempFiles=FALSE)

   setwd("D:/Leddiv/lilian/Bats/Disp_bar/MigClimTest")

   #Saving all species results in a single spreadsheet
  results <- read.table("MigClimTest_summary.txt", h=T, stringsAsFactors = FALSE)
  results[1,"simulName"] <- paste0 ( ext[i],"_rcp45")
  resu <- rbind (resu, results)

  # Reclassify raster
  # 0 : unsuitable;
  # 1 : climate refugia
  # 2 =< x =<29999 : colonized
  # x < 0 : locally extinct

  distr.sp <- raster("MigClimTest_raster.asc")
  #plot(distr.sp)

  m <- c(2, 29999, 2,    #Reclassify positives
         29999, 30000, 3)
  rclmat <- matrix(m, ncol=3, byrow=TRUE)
  rc <- reclassify(distr.sp, rclmat)

  s <- calc(rc, fun=function(x){ #Reclassify negatives
    x[x < 0] <- 4
    return(x)} )

  writeRaster(s, paste0("D:/Leddiv/lilian/Bats/Anthrop_maps/", ext[i],"_rcp45", ".tif"), format="GTiff", overwrite=T)
```

Note that the final map has 4 categories 0 : always unsuitable 1 : climate refugia (suitable in the present and in the future) 2 : potential colonization (became suitable and is accessible) 3 : dispersal limitation (became suitable but is unaccessible) 4 : non-analog climate (became unsuitable)

The same is applied to moderate habitat specialists, but the absence of tree is a weak barrier to them

```
### Moderate habitat specialists
# Barrier type = "weak"

for (i in 1: length(no_ext)) {

  setwd("D:/Leddiv/lilian/Bats/Disp_bar")

  a <- list.files(pattern = paste0(no_ext[i]))

  iniDist <- raster(paste0("D:/Leddiv/lilian/Bats/Mapas_Defaun/", no_ext[i], "_pres_PA.tif"))

  barr <- crop(barrier, extent(iniDist))
  barr <- as.data.frame(barr, xy=F)
  barr[is.na(barr)] <- 0

  iniDist <- as.data.frame(iniDist, xy=T)
  iniDist[is.na(iniDist)] <- 0

  nrow(barr) == nrow(iniDist)

  # rcp45
  setwd("D:/Leddiv/lilian/Bats/Disp_bar/")
  b <- stack(a)
  b45 <- raster::subset(b, grep("rcp45", names(b), value=T))
  b45@crs <- barrier@crs

  hsMap <- as.data.frame(b45)   # do not get coordinates

  nrow(hsMap) == nrow(iniDist)

  hsMap[is.na(hsMap)] <- 0      # all NAs converted to 0

   nrow(hsMap) == nrow(iniDist)
  nrow(barr) == nrow(iniDist) # TRUE Number of rows must match

  ### Dispersal

  MigClim.migrate(iniDist = iniDist,
                  hsMap = hsMap,
                  rcThreshold = 1, #artificial threshold, binarization was made during SDM
                  envChgSteps = ncol(hsMap), dispSteps = ncol(hsMap),
                  barrier = barr, barrierType = "weak", # Barrier is weak to moderate habitat specialists
                  replicateNb=3,
                  testMode=FALSE,
                  fullOutput=T, keepTempFiles=FALSE)

  setwd("D:/Leddiv/lilian/Bats/Disp_bar/MigClimTest")

  #Saving all species results in a single spreadsheet
  results <- read.table("MigClimTest_summary.txt", h=T, stringsAsFactors = FALSE)
  results[1,"simulName"] <- paste0 ( no_ext[i],"_rcp45")
  resu <- rbind (resu, results)

  distr.sp <- raster("MigClimTest_raster.asc")

  m <- c(2, 29999, 2,    #Reclassify positives
         29999, 30000, 3)
  rclmat <- matrix(m, ncol=3, byrow=TRUE)
  rc <- reclassify(distr.sp, rclmat)

  s <- calc(rc, fun=function(x){ #Reclassify negatives
    x[x < 0] <- 4
    return(x)} )

  writeRaster(s, paste0("D:/Leddiv/lilian/Bats/Anthrop_maps/", no_ext[i],"_rcp45", ".tif"), format="GTiff", overwrite=T)}
```

Yet for habitat generalists, the absence of trees is not considered a barrier.

```
### Habitat generalists
# Barrier file = empty

no.barr <- barrier
no.barr[no.barr==1] <- 0 # Transformed all barrier into non-barrier

for (i in 1: length(ge)) {

  setwd("D:/Leddiv/lilian/Bats/Disp_bar/")

  a <- list.files(pattern = paste0(ge[i]))
  iniDist <- raster(paste0("D:/Leddiv/lilian/Bats/Mapas_Defaun/", ge[i], "_pres_PA.tif"))
  barr <- crop(no.barr, extent(iniDist))

  iniDist <- as.data.frame(iniDist, xy=T)
  iniDist[is.na(iniDist)] <- 0

  barr <- as.data.frame(barr, xy=F)
  barr[is.na(barr)] <- 0

  # rcp45
  setwd("D:/Leddiv/lilian/Bats/Disp_bar/")
  b <- stack(a)
  b45 <- raster::subset(b, grep("rcp45", names(b), value=T))

  hsMap <- as.data.frame(b45)   # do not get coordinates
  nrow(hsMap) == nrow(iniDist)
  hsMap[is.na(hsMap)] <- 0      # all NAs converted to 0

  ### Dispersal

   MigClim.migrate(iniDist = iniDist,
                  hsMap = hsMap,
                  rcThreshold = 1, #artificial threshold, binarization was made during SDM
                  envChgSteps = ncol(hsMap), dispSteps = ncol(hsMap),
                  barrier = barr, #Barrier file is empty, but required to run simulations
                  replicateNb=3,
                  testMode=FALSE,
                  fullOutput=T, keepTempFiles=FALSE)

  setwd("D:/Leddiv/lilian/Bats/Disp_bar/MigClimTest")

  #Saving all species results in a single spreadsheet
  results <- read.table("MigClimTest_summary.txt", h=T, stringsAsFactors = FALSE)
  results[1,"simulName"] <- paste0 ( ge[i],"_rcp45")
  resu <- rbind (resu, results)

  distr.sp <- raster("MigClimTest_raster.asc")
  m <- c(2, 29999, 2,    #Reclassify positives
         29999, 30000, 3)
  rclmat <- matrix(m, ncol=3, byrow=TRUE)
  rc <- reclassify(distr.sp, rclmat)

  s <- calc(rc, fun=function(x){ #Reclassify negatives
    x[x < 0] <- 4
    return(x)} )

  writeRaster(s, paste0("D:/Leddiv/lilian/Bats/Anthrop_maps/", ge[i],"_rcp45", ".tif"), format="GTiff", overwrite=T)

  }
```
